# Species-specific responses to paleoclimatic changes and landscape barriers drive contrasting phylogeography of co-distributed lemur species in northeastern Madagascar

**DOI:** 10.1101/2025.06.11.658897

**Authors:** Tobias van Elst, Dominik Schüßler, Stephan M. Rafamantanantsoa, Tahiriniaina Radriarimanga, Naina R. Rabemananjara, David W. Rasolofoson, R. Doménico Randimbiharinirina, Paul A. Hohenlohe, Ute Radespiel

## Abstract

Madagascar is a megadiverse island with exceptionally high levels of endemism, which resulted mainly from allopatric speciation promoted by the island’s complex physical geography and paleoclimatic cycles. Northeastern Madagascar is uniquely suited to test the relative importance of river barriers, topography and climatic conditions for lemur genetic differentiation. Based on restriction site-associated DNA sequences, we inferred phylogeny, population structure, genetic diversity and rates of gene flow in four mouse lemur (genus *Microcebus*) and two woolly lemur (genus *Avahi*) species. In addition, we employed isolation-by-resistance models to test the importance of topography, river barriers, climate and forest cover in restricting intraspecific gene flow. Our results show that significant differences in genetic diversity and connectivity can be explained by varying responses to landscape features and species-specific phylogeographic histories. Rivers present a general barrier to gene flow, and dispersal between inter-river systems is predominantly mediated through high-elevation headwater regions. While this led to high connectivity and genetic diversity in *M. lehilahytsara* and *A. laniger*, gene flow among *M. jonahi* populations is limited by low climatic niche suitability at higher elevations. In addition, topographic complexity promotes connectivity among *Microcebus* populations, potentially by buffering impacts of seasonal or historic changes in climatic conditions. Finally, the more restricted distributions of *M. macarthurii*, *A. mooreorum* and, to some extent, *M. simmonsi* likely resulted from refugial dynamics and sea level fluctuations which led to geographic isolation, microendemism and secondary contact zones. Our findings generated informed hypotheses regarding the colonization history of the studied species while also having important implications for their conservation.

## 1 Introduction

The mechanisms and factors underlying the diversification of species are of central importance to evolutionary biology. Speciation typically results from a restriction of gene flow between populations, causing differentiation due to genetic drift and selection (Coyne & Orr, 2004; Mayr, 1942). The spatial separation of populations by geographic barriers such as water bodies or mountain ranges is considered the predominant mode of speciation in animals (allopatric speciation; Coyne & Orr, 2004). However, restriction of gene flow can also occur more gradually. For instance, isolation-by-distance explains population structure in many species (Sexton et al., 2014). Dispersal can similarly be limited through (continuous) landscape features other than distance (isolation-by-resistance; IBR; McRae et al., 2008). Divergence along such ecogeographic gradients (parapatric speciation) is well documented (e.g., Fenker et al., 2021; Jaynes et al., 2022; Myers et al., 2019).

Madagascar is one of the most biodiverse places on Earth, encompassing ca. 5% of the world’s described terrestrial species (Goodman, 2022). Most of this diversity is endemic to the island (e.g., ca. 90% of terrestrial and freshwater vertebrates), with high spatial species turnover (microendemism; Antonelli et al., 2022; Vences et al., 2009; Wilmé et al., 2006). Madagascar’s patterns of biodiversity have often been explained by allopatric and parapatric speciation resulting from the island’s complex physical geography (e.g., mountain and river barriers; e.g., Goodman & Ganzhorn, 2004; Martin, 1972; Schüßler et al., 2025) and changes in paleoclimatic conditions leading to cycles of isolation to glacial refugia and subsequent range expansion, particularly during the Quaternary (e.g., watershed model; Mercier & Wilmé, 2013; Wilmé et al., 2006; reviewed in Brown et al., 2014; Vences et al., 2009). Former glacial refugia are expected to harbor populations with comparably high genetic diversity, which decreases in the direction of postglacial range expansion (e.g., due to founder effects; Hewitt, 1996, 1999). Paleoclimatic dynamics could also have led to the fragmentation and divergence of populations due to sea level changes (sea levels at glacial maxima were up to 120 m lower than today; (Bintanja et al., 2005). The potential of ecogeographic barriers to limit gene flow and promote speciation ultimately depends on species-specific traits like habitat preference, ecological flexibility and dispersal ability, among others. Gaining a deeper understanding of diversification processes in Malagasy biota and their dependence on such traits will not only guide more effective conservation efforts, which are urgently required due to increasing overexploitation and associated habitat loss (Ralimanana et al., 2022), but will also generally shed light on the mechanisms underlying speciation and evolutionary radiations.

We selected a region in northeastern Madagascar (hereafter referred to as study region) located between Marojejy National Park (NP) and Betampona Special Nature Reserve (SNR; Supplementary Figs. S1, S2; Poelstra et al., 2021) that is well suited to empirically compare the relative importance and interplay of diversification factors in lemurs, a radiation of more than 100 species across five extant families (Mittermeier et al., 2023). The region harbors some of the largest remaining continuous humid forests (both high- and lowland) of the island, and anthropogenic habitat loss and fragmentation began only relatively recently there (Vieilledent et al., 2018). It is characterized by an elevational cline from the Central Highlands towards the east coast, with rivers of different size separating the region latitudinally into 17 inter-river systems (IRSs). The basin of one river, the Antainambalana River, was previously hypothesized to have served as a retreat-dispersion watershed for species during Pleistocene climatic fluctuations (Mercier & Wilmé, 2013; Wilmé et al., 2006). In addition, the island Sainte Marie (Nosy Boraha) lies on the continental shelf about 10 km away from the mainland and was likely connected to it during paleoclimatic sea level fluctuations (Bintanja et al., 2005; Rohling et al., 2022).

Almost one fourth of all described lemur species can be found in the study region (Mittermeier et al., 2023). This includes four of the 19 currently recognized mouse lemur species (genus *Microcebus*; *M. jonahi, M. macarthurii, M. lehilahytsara* and *M. simmonsi*; Poelstra et al., 2021; Schüßler, Blanco, et al., 2020; Tiley et al., 2022; van Elst et al., 2025) and two of the nine described woolly lemur species (genus *Avahi*; *A. laniger* and *A. mooreorum*; Lei et al., 2008; Zaramody et al., 2006). In addition, Louis and Lei (2016) hypothesized the presence of another yet undescribed *Microcebus* species (*M.* sp. #2) on the Masoala peninsula based on differentiation in mitochondrial genome sequences. Ecological and genetic work in the study region (Andriantompohavana et al., 2007; Poelstra et al., 2021; Schüßler, Blanco, et al., 2020; Tiley et al., 2022; Zaramody et al., 2006) indicated that lemur species differ greatly in their distributional areas and patterns of diversity, with *M. lehilahytsara* and *A. laniger* encompassing much larger distributions than other species in their genera. Due to their interspecific focus, however, the studies relied on few sampling sites per species, leaving most IRSs unsampled. Consequently, the intraspecific diversity, population structure and particularly the determinants of population differentiation of species in the region remain open questions. In addition, although riverine barriers, elevation and refugial dynamics during paleoclimatic oscillations have been suggested as major drivers of diversification and demographic history in *Microcebus* species (Martin, 1972; Olivieri et al., 2007; Radespiel et al., 2012; Teixeira, Montade, et al., 2021; Teixeira, Salmona, et al., 2021; van Elst et al., 2023; Weisrock et al., 2010; Wilmé et al., 2006; Yoder et al., 2016) the combined impact of such landscape variables on genetic differentiation has never been tested with model-based approaches in this genus, and only rarely in Malagasy mammals or primates more generally (e.g., Baden et al. 2019). The role of these factors in promoting diversification processes in the genus *Avahi* remains completely unknown.

Here, we employed comparative analyses of population structure, gene flow and genetic diversity based on restriction-associated DNA (RAD) sequencing across an extensive sample set to address these open questions in *Microcebus* and *Avahi* species in the study region. In addition, we performed isolation-by-resistance modelling to identify the relative role of topography, rivers, climate and vegetation in explaining their population genetic differentiation.

## 2 Material and methods

### 2.1 Sampling

We aimed to cover the entire distributions of the six species in the study region in our sampling by including high- and lowland sites in each IRS if possible (Figs. S1, S2). Between 2006 and 2022, we collected a total of 269 and 34 individual ear biopsies at 33 and 11 different localities following Radespiel et al. (2008) and Hokan et al. (2017) for the genera *Microcebus* and *Avahi*, respectively (Table S1). Our sampling significantly improves knowledge on the distributions of all species studied (see Supplementary results). Ear biopsies of four *A. occidentalis* outgroup individuals were also included. We complemented our data set with published sequences of 104 *Microcebus* individuals, including *M. lehilahytsara* populations south of the study region and *M. murinus* outgroups (Poelstra et al., 2021; Tiley et al., 2022; Supplementary Table S1).

### 2.2 RAD sequencing and genotyping

We generated single-digest restriction site associated DNA sequencing (RADseq) libraries using the enzyme *SbfI*. To do so, we extracted genomic DNA with a modified QIAGEN DNeasy Blood & Tissue Kit protocol (see Sgarlata et al., 2018). New RADseq libraries were prepared and paired-end sequenced to a target coverage of 15x following two different protocols (see Supplementary methods and Table S2 for details). Sequencing data were demultiplexed and adapters were trimmed with bcl2fastq v2.20 (Illumina, Inc.), discarding reads with final length smaller than 20 bases. As mentioned, we added published RADseq data of 104 *Microcebus* samples (Table S1; Poelstra et al., 2021; Tiley et al., 2022). For these, adapters were trimmed using Trimmomatic v0.39 (Bolger et al., 2014) with the following parameters: Leading: 3, Trailing: 3, Slidingwindow: 4:15, Minlen: 60.

Trimmed reads of *Microcebus* individuals were aligned against the *M. murinu*s reference genome Mmur 3.0 (Larsen et al., 2017) with BWA-MEM (Li & Durbin, 2009), retaining only those that mapped to autosomal scaffolds, had a mapping quality larger than 20 and were properly paired. Subsequently, we used the reference-based approach of Stacks v2.53 (Rochette et al., 2019) to call genotypes across individuals. For the genus *Avahi*, the *de novo* approach of Stacks was used to build a catalogue and call genotypes from trimmed reads of individuals with the parameters M = 2, m = 2 and n = 3 due to the lack of a reference genome. Parameter tuning was conducted following Paris et al. (2017). In this way, we created two genus-specific genotype call sets for phylogenetic inference and to compare estimates of genetic diversity as well as four (sister) species-specific sets (*M. jonahi* + *M. macarthurii*, *M. lehilahytsara*, *M. simmonsi* and *A. laniger* + *A. mooreorum*) to increase signal and enable more accurate filtering for population genomic analyses. All sets were filtered following FS6 filtering recommendations of O’Leary et al. (2018) with a modified threshold of 10% maximum missing data per site for the final filtering step, using VCFtools v0.1.17 (Danecek et al., 2011).

### 2.3 Phylogenetic inference

We performed maximum likelihood phylogenetic inference on the genus-specific genotype call sets, using the GTR+Γ model of sequence evolution in IQ-TREE v2.2.0 (Minh et al., 2020) while correcting for ascertainment bias. We estimated ultrafast bootstrap support and performed a Shimodaira-Hasegawa-like approximate likelihood ratio test (SH-aLRT; Guindon et al., 2010), using 1,000 replicates.

### 2.4 Population genetic structure and gene flow

We investigated population structure through ancestry inference and principal component analysis (PCA). Individual ancestries were inferred via maximum likelihood in ADMIXTURE v1.3.0 (Alexander et al., 2009), setting *K* from one to ten (six for *M. simmonsi*, as this was the number of sampled localities). Optimal *K* was estimated via cross validation. PCA was conducted in the R package ‘SNPRelate’ v1.32.2 (Zheng et al., 2012).

We inferred directional rates of gene flow between adjacent populations under a coalescent model with the R package ‘gene.flow.inference’ v0.0.0.9 (Lundgren & Ralph, 2019). As input to the model, we used the genetic distance *D_PS_* (= 1 – proportion of shared alleles; Bowcock et al., 1994) between populations. We ran the algorithm for a total of 8,000,000 generations with 12.5% pre-burn-in and 37.5% burn-in.

We also tested for introgression between *M. macarthurii* (IRS 5) and adjacent *M. jonahi* (IRS 6) as well as between *A. mooreorum* (IRSs 4 and 2/3, respectively) and adjacent *A. laniger* (IRS 5) with Patterson’s *D*-statistic (Patterson et al., 2012), as implemented in Dsuite v0.4 r38 (Malinsky et al., 2021). We used *M. lehilahytsara* and *A. occidentalis* as outgroup, respectively, and based the test on 15 randomly selected individuals per clade (or all individuals if less than 15 were available).

### 2.5 Drivers of population genetic structure

We first tested for the presence of isolation-by-distance in the four species *M. jonahi*, *M. lehilahytsara*, *M. simmonsi* and *A. laniger*, which can be considered the null model in landscape genetic analyses (Balkenhol et al., 2009; Jenkins et al., 2010). To do so, we performed a Mantel test based on Spearman’s rank correlation *r_s_* of genetic vs. geographic distances with 9,999 permutations in the R package ‘vegan’ v2.5-7 (Dixon, 2020). As a measure of genetic distance between populations, we again used *D_PS_* because it is less likely to be biased by unequal sample size and/or violations of assumptions than the traditionally used *F_ST_* (Peterman et al., 2014). Spatial deviations from a model of isolation-by-distance were visualized with Estimated Effective Migration Surfaces (EEMS; Petkova et al., 2015), which was run for 4,000,000 generations with a burn-in of 1,000,000 and using 1,000 demes. Unlike in inference of population structure, diversity and gene flow, *M. macarthurii* and *A. mooreorum* were not included alongside their sister species in these analyses given that intrinsic reproductive barriers could bias inference.

We then evaluated the relationship between five ecogeographic landscape variables potentially affecting lemur dispersal (elevation, landscape heterogeneity, rivers, climatic niche models, and forest cover) and genetic distances *D_PS_* between individuals through isolation-by-resistance models for the four species *M. jonahi*, *M. lehilahytsara*, *M. simmonsi* and *A. laniger* (see Supplementary methods and Table S1 for sample selection). Models were optimized with the gradient-based algorithm of the R package ‘radish’ v0.0.2 (Peterman & Pope, 2021), and their likelihood was inferred through multiple regression of landscape resistances with genetic distances, while allowing diagonal movement. Specifically, we used a log-linear conductance model and a maximum likelihood population effect parameterization for linear mixed-effect models (MLPE), which accounts for the non-independence of pairwise landscape and genetic distances, therefore providing an accurate approach for model inference based on individual distances (Clarke et al., 2002; Shirk et al., 2018; van Strien et al., 2014). We compared a total of 26 different models per species based on the Akaike Information Criterion (AIC), including the null model of IBD, all univariate models and multivariate models with up to three variables (Tables S4 – S7). Models with four or more predictors were not compared as many of these failed to reach convergence. Spatial rasters for the five predictor variables were created at a spatial resolution of 150 m as follows:

1. Elevation (Fig. S3) was encoded as a continuous variable based on the digital elevation model by GEBCO Compilation Group (2023).
2. Topographic complexity (Fig. S4) was encoded as a continuous variable by calculating landscape heterogeneity from the same digital elevation with the R package ‘rasterdiv’ v0.3.4 (Rocchini et al., 2021). In brief, this metric is based on a sliding window approach that assigns to each pixel an estimate of the variation among adjacent pixels.
3. Rivers (Fig. S5) were encoded as a continuous variable using flow accumulation, i.e., the amount of water that accumulates in a given area, as a proxy for size (O’Callaghan & Mark, 1984). Flow accumulation was calculated from the same digital elevation model with the R package ‘whitebox’ v2.2.0 (Lindsay, 2016). Rivers were padded to a width of two pixels to avoid diagonal traversal of rivers.
4. Climatic niche models (Figs. S6–S9) were inferred from spatially filtered occurrence records for each species (Table S3) and based on eight bioclimatic variables with ecological relevance for lemurs (see Supplementary methods for details).
5. The oldest available forest cover estimates of the region (i.e., from 1953) were obtained from Vieilledent et al. (2018) to approximate forest vegetation prior to human impact as accurately as possible (Fig. S10). Unpublished data by Jordi Salmona et al. indicate that these estimates (compared to more recent ones) best explain the population structure of *Propithecus coquereli* across its range.

All raster layers were cropped to the focal region of each species with a buffer of 35 km, centered around 0, and scaled by the standard deviation, using the R package ‘terra’ v 1.6.47 (Hijmans, 2022). Correlation coefficients among raster layers were below 0.7, based on a pixel subsample of 5% (Fig. S11).

### 2.6 Genetic diversity

To infer how genetic diversity is distributed across space, we estimated individual observed heterozygosities *H_O_* from the genus-specific genotype call sets with VCFtools. We quantified the relationships between *H_O_* and elevation, latitude and longitude in *M. jonahi*, *M. lehilahytsara*, *M. simmonsi* and *A. laniger* via Spearman’s rank correlation in R to identify potential glacial refugia (following Bagley et al., 2017). The correlation was not estimated in *M. macarthurii* and *A. mooreorum* due to the limited sample sizes. We also tested for significant differences between species of the same genus with a Kruskal-Wallis test and Dunn’s (*post hoc*) test with Bonferroni correction.

## 3 Results

### 3.1 RADseq and genotyping statistics

We sequenced an average of 9,342,093 (SD = 5,249,265) raw reads per sample, resulting in 80,560 (SD = 11,971) recovered RAD loci across *Microcebus* samples and 130,696 (SD = 27,188) loci across *Avahi* samples, with mean coverages of 10.86x (SD = 6.05x) and 9.98x (SD = 3.98x), respectively (Table S2). Two *Avahi* individuals from IRS 11 were excluded prior to genotyping due to low sequencing quality. While the genus-specific SNP sets included 157,306 (*Microcebus*) and 27,100 (*Avahi*) filtered variant sites, the (sister) species-specific SNP sets comprised 45,865 (*M. jonahi + M. macarthurii*), 112,138 (*M. lehilahytsara*), 25,838 (*M. simmonsi*), and 28,965 (*A. laniger* + *A. mooreorum*) variants after filtering. Proportions of missing data are given in Table S2.

### 3.2 Phylogenetic inference

*M. macarthurii* (IRS 5) and *M. jonahi* (IRSs 7–14) were recovered as reciprocally monophyletic (Figs. 1, S12). The deepest split in the *M. jonahi* phylogeny coincided with the Simianona River. The lineage north of this river formed two well-supported clades, one comprising populations from IRSs 6–8 and one those from IRSs 9–11 (separated by the Fahambahy River). Similarly, the well supported southern lineage exhibited a split between populations from IRSs 12 and 13/14, respectively (separated by the Marimbona River). More recent divergences were difficult to resolve, and support values were lower.

**Figure 1:**
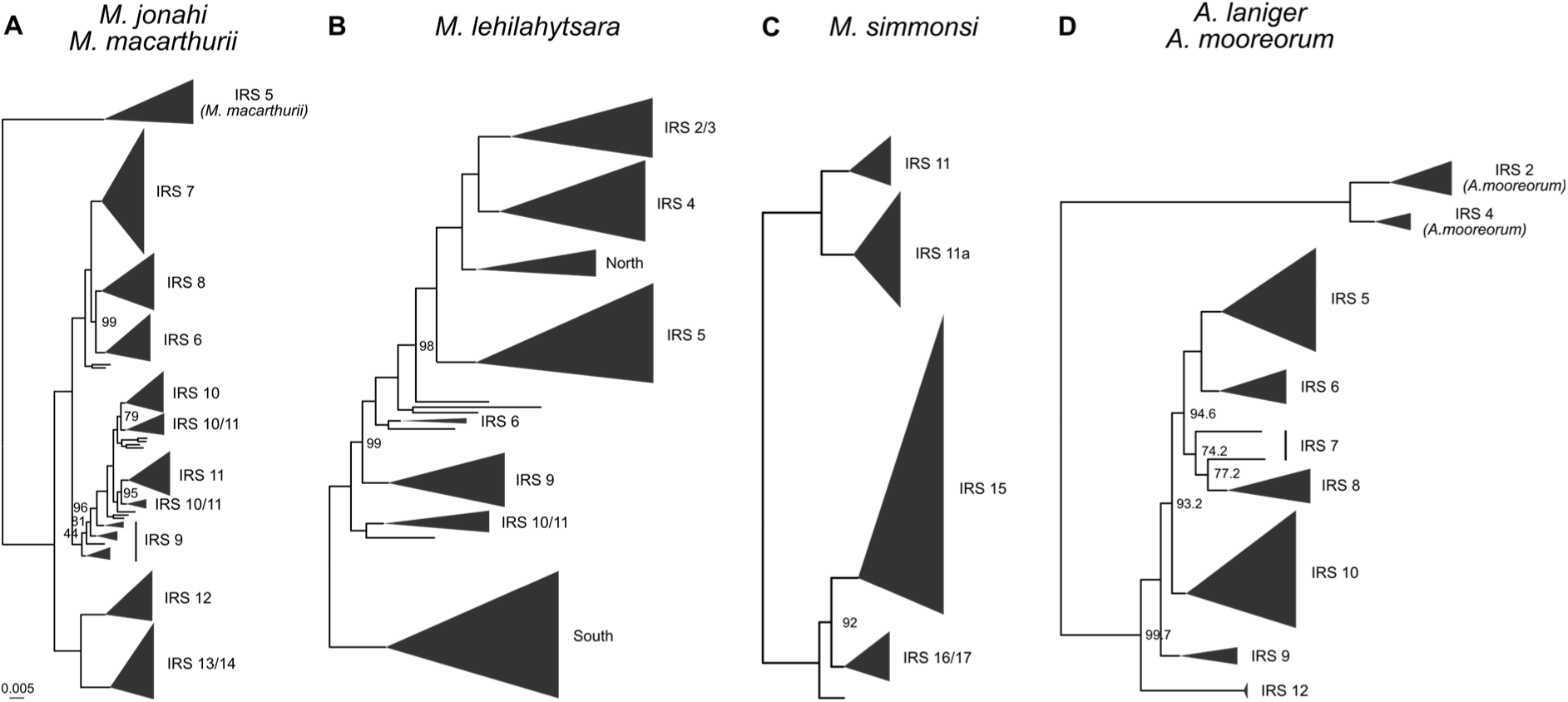
Maximum likelihood phylogenies inferred with IQ-TREE for *M. jonahi/M. macarthurii* (**A**), *M. lehilahytsara* (**B**), *M. simmonsi* (**C**) and *A. laniger*/*A. mooreorum* (**D**). Tip labels denote inter-river systems (IRSs). “North” and “South” correspond to *M. lehilahytsara* populations north (Anjanaharibe-Sud Special Reserve and Marojejy National Park) and south (see Tiley et al. 2022) of the study region, respectively. Triangles represent collapsed tips proportional to sample size. Node labels represent ultrafast bootstrap support, given only for nodes between major clades and if below 100. Scale is substitutions per site. Detailed phylogenies for each species are given in Figs. S12–S15.

As already shown in Tiley et al. (2022), *M. lehilahytsara* populations from central eastern and northeastern Madagascar formed two distinct clades (Figs. 1, S13). Among the northern clade, populations branched off successively in a south to north direction with high support, starting with those from IRSs 10 and 11, followed by IRS 9, 6, 5, 4 and 2/3. Highland samples from the far north of the study region (Anjanaharibe-Sud Special Reserve (SR) and Marojejy NP) clustered as sister to IRSs 2/3 and 4. Accordingly, individuals from Masoala (IRS 2/3), which have been hypothesized to represent a distinct species by Louis and Lei (2016), were nested deep inside the *M. lehilahytsara* phylogeny.

For *M. simmonsi*, we recovered one major clade formed by the northern populations in IRS 11 and on Île Ste. Marie (IRS 11a) and one comprising the southern populations (IRSs 15–17; Figs. 1, S14).

The divergence between *A. mooreorum* (IRSs 2 and 4) and *A. laniger* (IRSs 5 to 12) was well supported (Figs. 1, S15). Similar to *M. jonahi*, the Simianona River corresponded to the deepest split in the *A. laniger* phylogeny. Among populations north of this river, the population from IRS 9 represented the earliest divergence, followed by those from IRS 10 and a clade composed of sister lineages corresponding to IRSs 7 and 8 and IRSs 5 and 6, respectively.

### 3.3 Population genetic structure and gene flow

Cross validation of admixture analysis supported *K* = 6 as the best number of clusters for the set of *M. jonahi* and *M. macarthurii* individuals (Fig. S16). These clusters largely corresponded to phylogenetic clades and the regions between rivers with the highest-elevation headwaters (e.g., Rantabe, Simianona, and Marimbona Rivers) (Figs. 2A, S17). At *K* = 6, substantial admixture was only found in IRS 9 (i.e., between the populations north and south of it), which is formed by rivers with headwaters at lower elevations. The admixture results were also supported by PCA (Fig. S18), in which clusters were clearly separated and admixed individuals from IRS 9 fell in between populations north and south of them. The largest genetic distance among *M. jonahi* individuals was inferred between populations from IRSs 12– 14 and those north of them (clearly separated along PC1 in Fig. S18), the latter of which rather forming a gradient in genetic distance (along PC2). This gradient and the admixture in IRS 9 were congruent with the fact that the highest rates of gene flow were found in the region comprised by IRSs 6–11 (Fig. 2A; Table S8). Moreover, rates of gene flow from highland into lowland regions were generally higher than vice versa, and gene flow between highland regions was higher than that between lowland regions when considering the same adjacent IRSs. Notably, we also inferred gene flow between *M. macarthurii* and adjacent *M. jonahi* populations, which was supported by a significant test for introgression via Patterson’s *D*-statistic (*D* = 0.26, *Z*-score = 16.84, *p* < 0.0001; Fig. S19).

**Figure 2:**
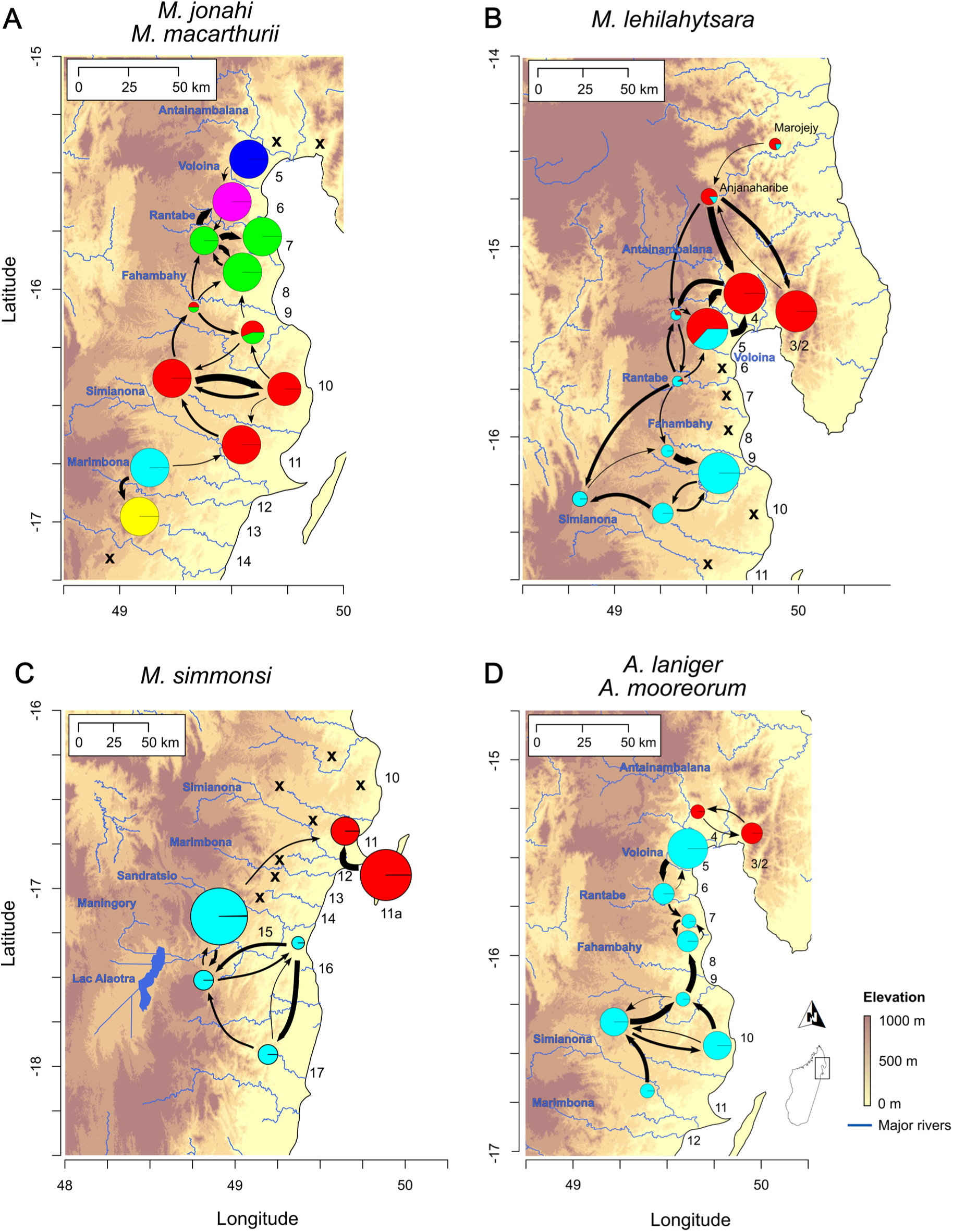
Population structure of *M. jonahi/M. macarthurii* (**A**), *M. lehilahytsara* (**B**), *M. simmonsi* (**C**) and *A. laniger*/*A. mooreorum* (**D**) in the study region in northeastern Madagascar. Colored pie charts show admixture proportions of sampled populations based on the best number of clusters (*K* = 6 for *M.jonahi*/*M. macarthurii*; *K* = 2 for the remaining taxa). In panel A and D, the blue and red pie charts corresponds to *M. macarthurii* and *A. mooreorum*, respectively. Pie chart size is proportional to sample size. Arrows indicate major gene flow between adjacent populations. Arrow width is proportional to relative gene flow rates. Please note that arrows represent gene flow but not necessarily migration routes (e.g., across a river). Crosses indicate absence records. Numbers denote inter-river systems. Blue labels denote names of rivers and Lac Alaotra.

In *M. lehilahytsara*, the best number of clusters was *K* = 2 (Fig. S16), indicating admixed ancestry of populations in IRSs 5 and 6 as well as in the far north of the study region, which represents the highlands of IRS 4 (Fig. 2B). Increasing *K* largely resulted in clusters corresponding to single IRSs (Fig. S20). The gradient-like pattern of admixed ancestry at *K* = 2 mirrored the successive branching recovered in phylogenetic inference and was also supported by PCA (Fig. S18). High levels of gene flow were found between populations throughout the entire region (Fig. 2B; Table S9), including those on opposite sides of the large Antainambalana River. Similar to *M. jonahi*, gene flow from highland into lowland populations was on average higher than vice versa.

The population structure of *M. simmonsi* was characterized by the high differentiation between the northern populations in IRS 11 and on Île Ste. Marie (IRS 11a) and the southern ones in IRSs 15–17, which was already evident in the phylogeny. These two clades corresponded to two distinct clusters in the admixture analysis (best *K* = 2; Fig. S16), which did not show substantial admixed ancestry (Fig. 2C). This was also evident in all other tested *K* (Fig. S21). In addition, the PCA supported high differentiation of the two clusters on PC1 (Fig. S18), and gene flow was predominantly inferred within them (with the exception of IRS 15 to IRS 11; Fig. 2C, Table S10). Gene flow within the northern clade was inferred mainly from IRS 11a to IRS 11.

Finally, *K*=2 was inferred as the best number of clusters for *A. laniger* and *A. mooreorum* (Fig. S16), with each species corresponding to one cluster (Fig. 2D). Increasing *K* indicated admixed ancestry of *A. laniger* individuals in IRSs 7–9 (Fig. S22), and high rates of gene flow between neighboring IRSs were inferred throughout the species’ entire range in northeastern Madagascar (Fig. 2D). This was also supported by principal component analysis, where adjacent IRSs clustered closely together (Fig. S23). Moreover, the coalescent model inferred minor gene flow (Table S11) between *A. laniger* (IRS 5) and *A. mooreorum* (IRS 4), which was congruent with a slight but significant excess of shared alleles between *A. laniger* (IRS 5) and *A. mooreorum* inferred by Patterson’s *D*-statistic (IRS 2: *D* = 0.06, *Z*-score = 4.22, *p* < 0.0001; Fig. S24).

### 3.4 Drivers of population genetic structure

A significant pattern of IBD across the distributional range was found in *M. jonahi* (*r_s_* = 0.82, *p* = 0.0001) and *M. lehilahytsara* (*r_s_* = 0.58, *p* = 0.0002) but not *in M. simmonsi* (all populations: *r_s_* = 0.39, p = 0.12; only southern populations: *r_s_* = 0.39, p = 0.12) or *A. laniger* (*r_s_* = 0.02, p = 0.45; Fig. S30). Spatial deviations from a null model of IBD aligned with major rivers in *M. jonahi* (negative log migration), e.g., those separating IRSs 6 and 7, 9 and 10, 11 and 12, or 12 and 13 (Fig. 3A). Log migration rates within IRSs were largely positive. In *M. lehilahytsara*, log migration rates were positive throughout much of the study region (Fig. 3B). Negative estimates were found around the Antainambalana River, the rivers south of Marojejy NP (in the far north), and the coastal areas at the northern end of Antongil Bay. *M. simmonsi* showed negative log migration values between the northern clade (IRSs 11 and 11a) and the southern clade (IRS 15–17) and, to a smaller extent, between populations in IRS 15 and those in IRSs 16 and 17 (Fig. 3C). Finally, in *A. laniger*, negative log migration rates were found between IRSs 10/11 and 12 as well as around IRSs 7 and 8 (Fig. 3D).

**Figure 3:**
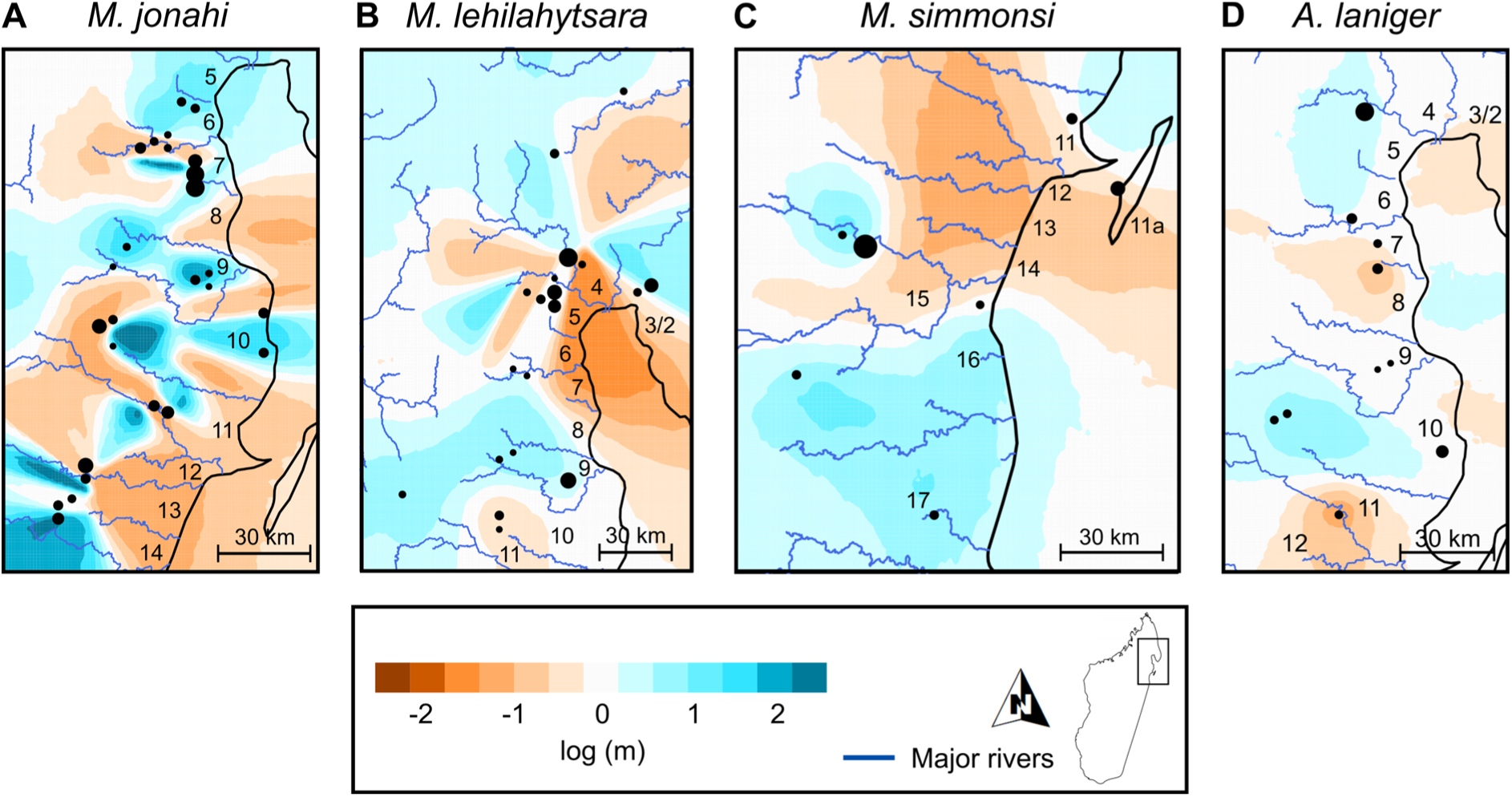
Estimated effective migrations surfaces (EEMS) for the four species *M. jonahi* (**A**)*, M. lehilahytsara* (**B**)*, M. simmonsi* (**C**) and *A. laniger* (**D**) in northeastern Madagascar. Effective migration is displayed on log_10_ scale. Numbers indicate inter-river systems.

All univariate isolation-by-resistance models performed better (measured by AIC) than the null model of isolation-by-distance in *M. jonahi* and *M. lehilahytsara* (except for elevation in *M. lehilahytsara*; Tables S4, S5). In addition, multivariate models generally had a better fit than univariate models. The best fitting model in *M. jonahi* retained flow accumulation, climatic niche suitability and landscape heterogeneity as significant predictors (Fig. 4A). While flow accumulation imposed a high negative effect on conductance, climatic niche suitability and landscape heterogeneity promoted conductance in *M. jonahi*. In *M. lehilahytsara*, flow accumulation (negative effect), landscape heterogeneity and forest cover (both with a positive effect) were significant predictors in the best fitting model (Fig. 4B). Conversely, no model performed significantly better than the null model in *M. simmonsi* and *A. laniger* (i.e., the null model had a ΔAIC < 2 compared to the best alternative model; Tables S6, S7).

**Figure 4:**
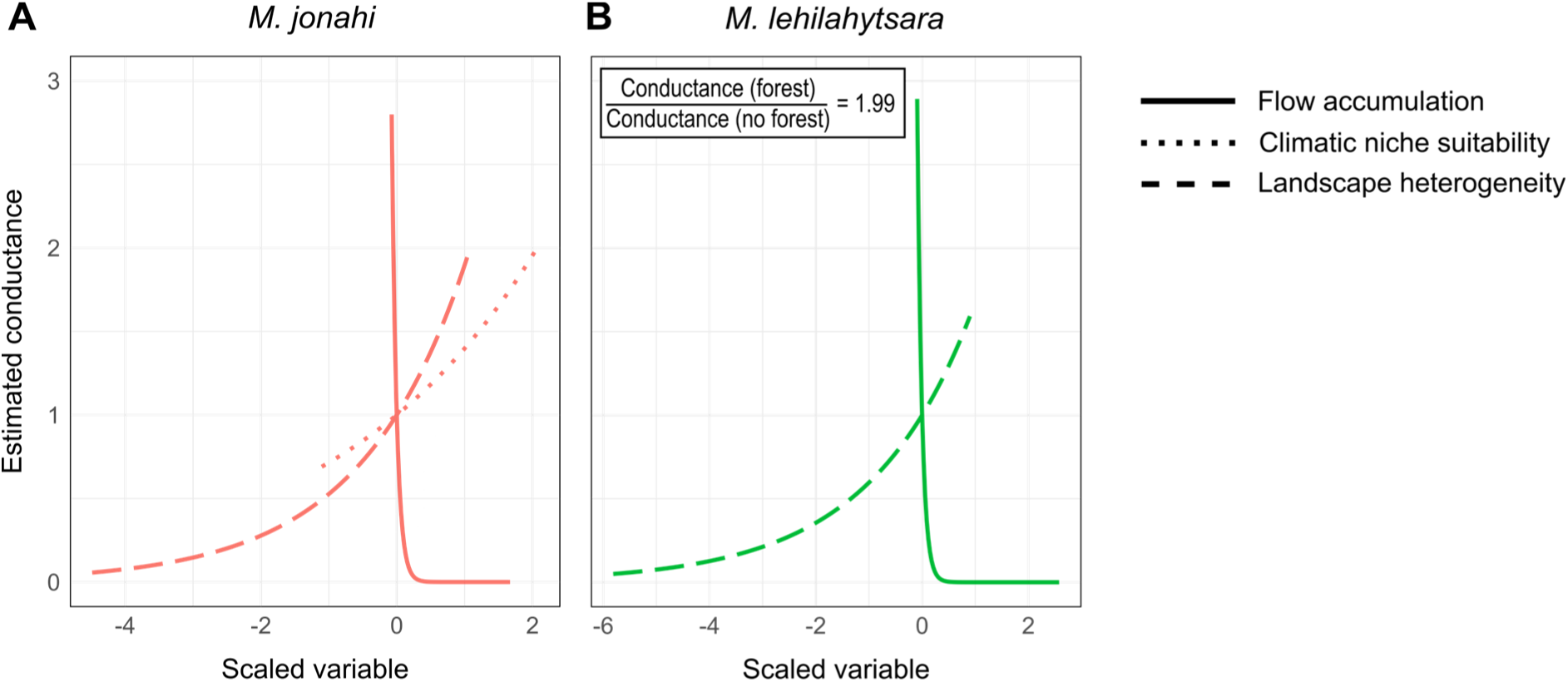
Estimated conductance associated with scaled values of predictor variables with significant effect in the best fitting models of isolation-by-resistance in *M. jonahi* (**A**; red) and *M. lehilahytsara* (**B**; green). The relative conductance of the forest feature (categorical variable) in comparison to its absence is given in the inlet for *M. lehilahytsara*.

### 3.5 Genetic diversity

There were significant differences in *H_O_* among species in the study region (Kruskal-Wallis test: *H* = 319.65, *df* = 8, *p* < 0.0001; Fig. S25, Tables S12, S13). Specifically, *H_O_* was significantly higher in *Avahi* than in *Microcebus* species, except for *M. lehilahytsara.* Within the genus *Microcebus*, mean *H_O_* was approximately three times higher in *M. lehilahytsara* than in the other species (Dunn’s test: to *M. jonahi*: *z* = 11.44, *p* < 0.0001; to *M. macarthurii*: *z* = −9.10, *p* < 0.0001; to *M. simmonsi*: *z* = −11.63, *p* < 0.0001). In addition, *M. jonahi* had significantly higher *H_O_* than *M. macarthurii* (*z* = −3.83, *p* = 0.005) and *M. simmonsi* (*z* = −4.37, *p* < 0.0001). No significant differences were found in *H_O_* between *Avahi* species, probably due to limited sampling size. However, mean *H_O_* of *A. laniger* was approximately 1.5 times higher than that of *A. mooreorum*.

The correlations between *H_O_* and latitude, longitude and elevation were significant in *M. jonahi* and *M. simmonsi* (Figs. S26, S27), but not in *M. lehilahytsara* (Fig. S28). In *M. jonahi*, *H_O_* showed a strong positive correlation with latitude (higher *H_O_* was found at more northern regions; *r_s_* = 0.86, *p* < 0.0001). The relationships with respect to longitude (*r_s_* = 0.48, *p* < 0.0001) and elevation (*r_s_* = −0.20, *p* = 0.0083) were also significant but weaker. In contrast, in *M. simmonsi*, lower *H_O_* was found at more northern and eastern sampling sites (*r_s_* = −0.39, *p* = 0.0185 and *r_s_* = −0.56, *p* = 0.0004 for latitude and longitude, respectively) and at higher elevations (*r_s_* = 0.47, *p* = 0.004). This pattern was largely driven by the fact that the northern *M. simmonsi* clade (IRS 11 and 11a) exhibited the highest latitude and longitude and the lowest elevation and had lower mean *H_O_* estimates than the southern clade. In *A. laniger*, *H_O_* was only correlated significantly with latitude (*r_s_* = 0.57, *p* = 0.0019; Fig. S29). It is important to note that, due to the topographical nature of the study region, longitude and elevation were negatively correlated in all four species.

## 4 Discussion

### 4.1 Population structure and its determinants

We reconstructed the population genetic structure of four *Microcebus* and two *Avahi* species in northeastern Madagascar based on extensive sampling and examined the effects of various landscape features on gene flow. Because human pressure and associated habitat degradation in the region started relatively recently (Ralimanana et al., 2022; Schüßler, Mantilla-Contreras, et al., 2020; Vieilledent et al., 2018), we assume that inferred genomic signatures are predominantly the result of non-anthropogenic factors such as geographic features and historic climatic processes.

We detected substantial population structure in *M. jonahi*, with genetic clusters largely corresponding to different IRSs (Fig. 2A). Congruently, effective migration surfaces (Fig. 3A) and IBR models (Fig. 4) revealed that rivers are important barriers to gene flow in this species, as suggested previously (Martin, 1972; van Elst et al., 2023). Consequently, migration corridors between IRSs are only given in headwater regions where rivers become small enough to be bridged by vegetation and crossed by arboreal animals (Goodman & Ganzhorn, 2004; Wilmé et al., 2006). However, these routes may be constrained for *M. jonahi* given that climatic niche suitability was a significant predictor of connectivity (Fig. 4) and declines at higher elevations under current climatic conditions (Fig. S11). Consistent with this, admixture and high rates of gene flow were predominantly found between IRSs where vegetation bridges across rivers exist today at these or lower elevations (IRSs 6–11; Fig. 2A), whereas the highest differentiation (between populations in IRS 12–14 and those to the north) coincided with the Simianona River, which is bridged by vegetation only at ca. 1,000 m a.s.l. Similar patterns of isolation by river barriers and elevational tolerance in Madagascar’s humid forests were shown for *M. gerpi* (van Elst et al., 2023).

Our findings also indicated that, while elevation presents a good proxy, it cannot explain dispersal limitations around river headwaters alone because it was only moderately correlated with climatic niche suitability (Fig. S11) and did not appear in the best-fitting IBR model for *M. jonahi* (Table S4). Rather, more intricate (micro-)climatic and ecological conditions that cannot be easily predicted by elevation alone seem to limit the species’ dispersal ability. This hypothesis is supported by the positive relationship of topographic complexity (i.e., landscape heterogeneity) and genetic connectivity in *M. jonahi* (and *M. lehilahytsara*; Fig. 4). Topographically complex regions may offer a range of relatively stable microclimates due to protective sheltering effects in connected habitat pockets, that may be used as migratory corridors. These advantageous ecological settings may facilitate the buffering of diurnal, seasonal and unpredictable interannual and historic climatic fluctuations (e.g., in temperature or humidity;Byrne et al., 2022; John et al., 2024). Such a buffering effect is expected to be particularly relevant in Madagascar given its highly seasonal climate (Goodman, 2022), interannual variation in climatic conditions (Dewar & Richard, 2007) and recurring natural disasters like cyclones (Donque, 1975) and droughts (e.g., Gould et al., 1999; Virah-Sawmy et al., 2010), the latter of which can also severely impact the humid rainforests of the island (Ranomafana NP, unpubl. records). It has already been suggested that these factors play a role in explaining the idiosyncratic life histories of lemurs compared to other primates (Dewar & Richard, 2007; Wright, 1999). Accordingly, topographically complex regions may serve as retreat areas for mouse lemurs and other taxa, potentially promoting higher population sizes, higher genetic diversity and dispersal. Such regions may also be associated with less human pressure (e.g., habitat loss and fragmentation) as it should be more difficult to access and cultivate topographically complex regions (Schüßler, Mantilla-Contreras, et al., 2020). However, we did not find high correlations between landscape heterogeneity and proxies for anthropogenic pressure like forest cover (or a lack thereof; Fig. S11) or human population density (which was explored for approximately 75% of the study region; D. Schüßler, unpubl. data).

While Poelstra et al. (2021) detected genetic differentiation between *M. jonahi* and *M. macarthurii*, they were not able to exclude the possibility that sampled populations represented opposite ends of a cline in genetic variation due to a lack of data for IRSs 6–8. We filled this sampling gap and showed that the two species are reciprocally monophyletic (Fig. 1A), with high genetic distances (even at small geographic distances; Fig. S30) and significant differences in genetic diversity (Fig. S25, Table S13). Accordingly, our findings confirm the species level distinction of the two taxa originally suggested by Poelstra et al. (2021) and Schüßler, Blanco, et al. (2020). Like several other *Microcebus* species (e.g., *M. berthae, M. sambiranensis*; see van Elst et al., 2025), *M. macarthurii* seems to be an extreme microendemic with a restricted range in the lowland forests of IRS 5 and on the island of Nosy Mangabe (not sampled; Louis & Lei, 2016), explaining its low genetic diversity. Notably, we inferred low levels of gene flow between the two species (Fig. 2A), which was also supported by low levels of admixture at several *K* and a significant test for introgression (Figs. S17, S19). This mirrors previous findings of mitochondrial introgression from *M. jonahi* into *M. macarthurii* by Poelstra et al. (2021) and Radespiel et al. (2008). How recent this signal is, remains to be explored.

In contrast to *M. jonahi*, *M. lehilahytsara* populations were characterized by high rates of gene flow between IRSs (Fig. 2B) and high genetic diversity throughout the entire range of the species in northeastern Madagascar (Fig. S28). This included the population on the Masoala peninsula (IRS 2), which was hypothesized to harbor a distinct *Microcebus* species (i.e., *M.* sp. #2) by Louis and Lei (2016). Our results suggest that *M.* sp. #2 is not a separate species but part of a large-scale geographic cline in genetic differentiation across more than 600 km, which includes *M. lehilahytsara* populations from central eastern Madagascar (e.g., Tsinjoarivo) and extends through IRSs 5–11 up to Anjanaharibe-Sud SR and Marojejy NP, including the former *M. mittermeieri* (Poelstra et al., 2021; Tiley et al., 2022). This finding highlights the importance of complete spatial sampling for characterizing intraspecific (genetic) variation when inferring taxonomic relationships. While rivers were also inferred as strict barriers to gene flow in *M. lehilahytsara* (Fig. 4B), high rates of gene flow among adjacent highland populations, together with gene flow predominantly directed from high-into lowlands (Fig. 2C), suggest that the species’ high connectivity and genetic diversity are likely maintained by extensive migration through highland regions. This enables the crossing of rivers at headwaters (Goodman & Ganzhorn, 2004; Wilmé et al., 2006), including even major river systems originating at high elevations such as the Antainambalana River. This hypothesis is further supported by a lack of elevation and climatic niche as significant predictors in the best fitting IBR model (Table S5), indicating that this species possesses high habitat flexibility (see also Andriambeloson et al., 2021; Schüßler, van Elst, et al., 2025). This is potentially due to physiological or behavioral adaptations to environmental challenges associated with higher elevations (e.g., the use of prolonged torpor to buffer against unfavorable environmental conditions; Blanco et al., 2017).

While population structure in *M. jonahi* and *M. lehilahytsara* was better explained by models of IBR than by the null model of IBD, this was not the case for *M. simmonsi*, with low variation in AIC (Table S6). However, to avoid biases from disjunct sampling and high differentiation of the northern and southern lineages, we based IBR models in this species only on samples from the southern part of the species distribution (four populations; IRS 15–17). This limited sampling for *M. simmonsi* may thus not have provided sufficient power to detect signatures of genetic differentiation promoted by landscape features. Nevertheless, the high inferred connectivity even across river barriers and low differentiation among southern populations (Fig. 2C) suggests that this species has a relatively high elevational tolerance. For instance, gene flow between the highlands of IRS 15 and IRS 16 may be mediated by dispersal through the marshes of Lac Alaotra, where mouse lemurs have already been reported, even though their species identity remains to be confirmed (Garbutt, 2022).

Similar to *M. lehilahytsara*, *Avahi laniger* was characterized by only weak population structure and high rates of gene flow throughout the sampled region (Fig. 2D), although phylogenetic inference (Fig. 1D) and admixture proportions at higher *K* (Fig. S22) indicated that some rivers correlate with genetic structure (e.g., the Rantabe and Simianona Rivers separating IRSs 6/7 and 11/12, respectively, as also observed in *M. jonahi*). This suggests that *A. laniger* disperses effectively across river barriers around headwater regions similar to *M. lehilahytsara* (we consistently sighted *Avahi* individuals in headwater regions of IRSs 13-15 but were unable to collect samples at those sites). However, we acknowledge that the lack of significant IBR signals (e.g., barrier function of rivers) may also partly be the result of a lack of power due to limited sampling. Notably, we showed that *A. laniger* does not occur in IRSs 2/3 and 4, as our genomic data identified individuals there as *A. mooreorum*, supporting the substantial barrier function of the Antainambalana River. The supposed presence of *A. laniger* in Anjanaharibe-Sud SR (Lei et al., 2008) suggests, however, that the species does disperse around the river’s headwaters but has not been able to colonize the Masoala peninsula, like *Indri indri* (Nunziata et al., 2016). Its further expansion towards Masoala may have been precluded by interspecific competition with *A. mooreorum*. The identified introgression signal between northern *A. laniger* and *A. mooreorum* indicates the presence of a secondary contact zone between the two sister species (a phylogeographic scenario for their initial split is illustrated in the next section), which is likely located between IRS 4 and Anjanaharibe-Sud SR. Congruent with the high connectivity among *A. laniger* populations, the species exhibited relatively high observed heterozygosity estimates throughout its range, which were approximately 1.5- and 3-fold higher than those found in *A. mooreorum* and *A. occidentalis*, respectively (Fig. S25, Table S13). It is important to note that heterozygosity estimates between the genera *Microcebus* and *Avahi* are not directly comparable, as different genotyping and filtering approaches were used (reference-based vs *de novo*, respectively). A reference-free approach may be more susceptible to genotyping errors, potentially leading to inflated estimates of heterozygosity.

### 4.2 Phylogeographic history and distribution

We can only fully understand the population structure und distribution of species by investigating their phylogeographic history (Hewitt, 1999). According to coalescent models by Poelstra et al. (2021) and van Elst et al. (2025) the diversification of *M. jonahi* and *M. macarthurii* took place during the Quaternary glaciation cycles of the past 200,000 years. Given *M. macarthurii*’s current distribution in IRS 5, north of *M. jonahi*, and the gradual decline in genetic diversity towards the southern end of *M. jonahi*’s range (Fig. S26), we hypothesize that their common ancestor colonized the study region during this time from central northern highland refugium. While the Antainambalana River, with its comparably high-elevation source at ca. 1,450 m a.s.l. (Goodman & Ganzhorn, 2004), likely served as a northern dispersal barrier for *M. macarthurii*, colonization towards the south by ancestral *M. jonahi* may have been facilitated by the lower-elevation headwaters of rivers separating IRSs 6–11. The initial colonization of IRS 12–14 despite the high-elevation headwaters of the Simianona and Marimbona Rivers may be explained by more favorable (warmer) climatic conditions during past interglacial periods, possibly allowing *M. jonahi* to disperse via higher elevations (Wilmé et al., 2006). Potentially, historic highland populations of *M. jonahi* may have subsequently lost favorable habitats with an increasing aridification during the LGM or have been outcompeted by the newly arriving *M. lehilahytsara* following their south to north colonization of the same region ca. 100 ka ago (Tiley et al., 2022). However, given that *M. jonahi* is significantly larger than *M. lehilahytsara* (Schüßler et al., 2023) and that *M. lehilahytsara* seems to shift its ecological niche when living in sympatry with *M. jonahi* (Schüßler, van Elst, et al., 2025), the latter explanation is not very likely.

Intriguingly, the divergence of *M. macarthurii* and *M. jonahi* cannot be explained by landscape features investigated in this study, as the river separating their two ranges has low-elevation headwaters with high habitat suitability for *M. jonahi* (Fig. S6) and a continuous forest cover across both ranges (Fig. S10). Potentially, the divergence of the two species occurred due to allopatric speciation in separate refugia during drier periods. In this scenario, populations in IRS 5 (today’s *M. macarthurii*) may have been confined to the area around the Antainambalana River, whereas the ancestral *M. jonahi* may have contracted to another refugium further south, as the Voloina River likely ran dry due to its lower-elevation headwater (Mercier & Wilmé, 2013; Wilmé et al., 2006). Such climatic conditions and associated lower sea levels would have also allowed the colonization of Nosy Mangabe by *M. macarthurii*, which is located close to the mouth of the Antainambalana River (Schüßler, 2025). *M. jonahi*’s missing recolonization of IRS 5 could be due to competition with its sister species for critical resources, as has been hypothesized for *M. jonahi* and *M. simmonsi* in IRS 11 (Schüßler, van Elst, et al., 2025).

The wide distribution of *M. lehilahytsara* in central eastern Madagascar (Tiley et al., 2022) and the successive phylogenetic divergences of populations in different IRSs (Fig. 1B) suggest that this species colonized the IRSs of the study region through range expansion over highland migration corridors from the south towards the north (similar to the hypothesized expansion of *M. murinus* in western Madagascar; Blair et al., 2014; Schneider et al., 2010; but see Fauskee et al., 2024). This allowed *M. lehilahytsara* to disperse around riverine barriers, including the large Antainambalana River, eventually reaching the Masoala peninsula. Such a migration scenario is supported by the admixed ancestry of populations north of this river (in Anjanaharibe-Sud SR and Marojejy NP) at *K* = 2 and their intermediate position in the PCA (Fig. S18). It may have been facilitated by favorable dispersal conditions during the Last Interglacial. The exceptional colonization potential of *M. lehilahytsara* together with its ecological plasticity (Schüßler, van Elst, et al., 2025) helps to explain its extensive range along the highlands of Madagascar’s east coast. In fact, as the entire east coast is separated into IRSs, the potential for highland migration is likely one of the most significant predictors of eastern mouse lemur range sizes in this region (see also van Elst et al., 2023). Notably, we did not find *M. lehilahytsara* in lowland regions of IRSs 6–8, 10 and 11 (Fig. S1), which could be due to insufficient sampling or competitive exclusion by the larger-bodied sympatric species, *M. jonahi* and *M. simmonsi*, in their core habitats. However, both explanations seem unlikely. First, our absence records are based on 13–30 individuals from multiple sites and forest types per IRS (Table S1), which would have enabled the detection of *M. lehilahytsara* even if they preferred different microhabitats or forests. Second, *M. lehilahytsara* occurs sympatrically at lower elevations with *M. macarthurii* or *M. jonahi* in IRS 5 and 9, respectively (Fig. S1). Alternatively, it appears more likely that these absences are the result of historic colonization and local extinctions in lowlands during the LGM. The larger-bodied lowland mouse lemurs could have survived the coldest and driest conditions in all IRSs by a stronger reliance on heterothermy, which may not have been possible for small-bodied *M. lehilahytsara* (Schüßler, van Elst, et al., 2025). Subsequent recolonization could have been impeded by niche competition with *M. jonahi* and *M. simmonsi* (Schüßler, van Elst, et al., 2025).

The main question with respect to the phylogeography of *M. simmonsi* pertains to its disjunct distribution and the high differentiation between northern (IRSs 11 and 11a) and southern (IRSs 15–17) populations (Fig. 2C). The colonization of Île Ste. Marie (IRS 11a) and IRS 11 may have been enabled by sea level changes associated with glacial periods. Extrapolating river courses via flow accumulation from a digital elevation model (Schüßler, 2025) suggests that the rivers separating IRS 11 from the southern *M. simmonsi* populations converged into a single river when sea levels were about 50–100 m lower than today (e.g., during the LGM; Fig. S31). Accordingly, only one dispersal event across this river (e.g., through rafting or meander cutoff; Haffer 2008) would have been necessary to colonize the area of Île Ste. Marie from IRS 16 and, subsequently, IRS 11, which was likely not separated from the current island by any water body during these times (Fig. S31). The newly arrived *M. simmonsi* may not have been able to further expand its range despite elevational tolerance due to competition with *M. jonahi* or larger-bodied lemur species that already inhabited IRSs 11–14 (Schüßler, Blanco, et al., 2020; Thorén et al., 2011). If *M. simmonsi* is indeed absent from IRSs 12–14 and anthropogenic causes are excluded (e.g., Pastorini et al. 2003), such a dispersal scenario currently represents the most parsimonious explanation for the disjunct distribution of this species.

Even though direct divergence time estimates are not available for *A. laniger* and *A. mooreorum*, Herrera & Dávalos (2016) indicated that the diversification of the genera *Microcebus* and *Avahi* in eastern Madagascar occurred on a similar time scale. Therefore, the paleoclimatic dynamics that affected mouse lemurs (van Elst et al., 2023) may have also shaped the diversification of *Avahi* species. Specifically, we hypothesize that the ancestor of the two sister species studied here may have become separated in different refugia on opposite sides of the Antainambalana River during more arid conditions, when dispersal corridors in highland regions became unavailable. Given the inferred phylogeny (Fig. 1D), the lineage south of the Antainambalana likely underwent a southward retreat whereas the other lineage remained restricted to the region around the Masoala peninsula, leading to their divergence. The subsequent northward recolonization of the study region by *A. laniger* and its dispersal around the Antainambalana River led to the current distribution patterns and a potential secondary contact zone. The fact that the distribution of *A. mooreorum* remained relatively restricted could also be due to competition with its sister species, as hypothesized for *M. macarthurii* and *M. jonahi* in IRS 5 and *M. simmonsi* and *M. jonahi* in IRS 11. As observed heterozygosities were similarly high across the entire range of *A. laniger* (likely due to its high connectivity; Fig. S29), our data do not allow to identify a specific refugium and colonization route.

Ultimately, the diversification scenarios formulated here will need to be confirmed through more detailed characterizations of the three identified (secondary) contact zones (*M. macarthurii*/*M. jonahi*, *M. jonahi*/*M. simmonsi*, *A. laniger*/*A. mooreorum*) and demographic models which enable identifying the timing of divergence events as well as changes in gene flow and population sizes associated with potential range expansions (e.g., Teixeira, Montade, et al., 2021; Teixeira, Salmona, et al., 2021).

### 4.3 Conservation implications

Our comparative investigation of population structure and its determinants revealed significant differences in genetic diversity and connectivity among *Microcebus* and *Avahi* species in northeastern Madagascar. These differences can in part be explained by varying responses of the species and genera to different landscape features. Although rivers present barriers to gene flow in all species, connectivity among IRSs is maintained over highland and headwater migration corridors in different ways. While comparably high connectivity and genetic diversity are present in *M. lehilahytsara* and *A. laniger*, gene flow among *M. jonahi* populations from different IRSs seems to be constrained by low climatic niche suitability at higher elevations. Accordingly, the protection of forest habitat around headwaters, where vegetation (e.g., the canopy) allows arboreal species to cross river barriers, should be prioritized in conservation efforts. This is particularly relevant considering potential scenarios of climate change and associated migratory movements of species to track changing habitats, which are predicted to become prominent in northeastern Madagascar (Brown & Yoder, 2015). The extensive Makira Natural Park already protects headwater regions in the northern part of the distribution of *M. jonahi* (extending south until IRS 7), but its southern half, which harbors populations with the lowest genetic diversity (Fig. S26), and potential migration corridors remain largely unprotected (except for Mananara-Nord NP in the lowlands of IRS 10 and Ambatovaky SR in the highlands of IRS 12; (Ralimanana et al., 2022). Particular attention should also be given to an effective protection of lowland forest habitats of IRS 5 (Anjiahely) and the island of Nosy Mangabe as these seem to be the only regions harboring the microendemic *M. macarthurii*, which already shows low genetic diversity compared to the other studied *Microcebus* species (currently protected as Makira Natural Park and Nosy Mangabe NP). The isolated *M. simmonsi* populations in IRS 11 (Ambodiriana) and on Île Ste. Marie (IRS 11a) also deserve urgent conservation attention, as to date, no formal protection status has been given to these sites (Ralimanana et al., 2022). Finally, the potential function of topographically complex regions as ecological buffers during seasonal and stochastic variations in environmental and climatic conditions requires further ecological investigation, as this could make them priorities for conservation efforts, too.

## Author contributions

**T. van Elst**: Conceptualization (equal); data curation (lead); formal analysis (lead); funding acquisition (supporting); investigation (lead); methodology (lead); project administration (equal); software (lead); resources (equal); validation (lead); visualization (lead); writing – original draft (lead). **D. Schüßler:** Conceptualization (supporting); data curation (supporting); funding acquisition (supporting); project administration (equal); resources (equal); visualization (supporting); writing – original draft (supporting). **S. M. Rafamantanantsoa:** Project administration (supporting); resources (supporting). **T. Radriarimanga:** Project administration (supporting); resources (supporting). **N. R. Rabemananjara:** Project administration (supporting); resources (supporting). **D. W. Rasolofoson:** Project administration (supporting); resources (supporting). **R. D. Randimbiharinirina:** Project administration (supporting); resources (supporting). **P. A. Hohenlohe:** Investigation (supporting); methodology (supporting); resources (supporting). **U. Radespiel:** Conceptualization (equal); data curation (supporting); funding acquisition (lead); investigation (supporting); methodology (supporting); project administration (equal); resources (equal); supervision (lead); validation (supporting); writing – review and editing (supporting).

## Acknowledgements

We thank the Ministry of Environment and Sustainable Development (formerly the Ministry of Environment, Water and Forests) of the Malagasy Government and its local representatives for allowing us to conduct the field work under the permits 197/17/MEEF/SG/DGF/DSAP/SCB.Re, 169/19/MEDD/SG/DGEF/DGRNE, 349/21/MEDD/SG/DGGE/DAPRNE/SCBE.Re and 030/22/MEDD/SG/DGGE/DAPRNE/SCBE.Re. All field procedures adhered to Malagasy regulations, standards of the International Primatological Society (International Primatological Society, 2014) and the “proposal for ethical research conduct in Madagascar” (Wilmé et al., 2016). We are grateful to B. M. Raharivololona and GERP (Groupe d’Etude et de Recherche sur les Primates) for help with project coordination and infrastructure. We also thank A. N. Raharinirina, D. Rabesamihanta, R. Rakotondravony, S. T. Rovanirina, N. T. Andriamaroroka, R. Andriamalala, E. Rasolondraibe, and all local guides for their invaluable help during sample collection. Finally, we are grateful to Biosearch Technologies (LGC Genomics GmbH, Berlin) and B. Fartmann for RADseq library preparation and to J. Salmona for advice and feedback on the isolation-by-resistance analysis.

## Funding

This project was funded by the German Research Foundation (DFG Ra 502/23-1 to U. R.), the Society for Tropical Ecology, Houston Zoo, Inc., the Bauer Foundation (T237/22985/2012/kg to D. S.), the Zemplin Foundation (T0214/32083/2018/sm to D. S.) and through a compute project provided by the German National High Performance Computing Alliance NHR@Göttingen (nib00015 to U. R. and T. v. E.).

## Conflict of interest

The authors declare no competing interests.

## Data availability statement

All new sequencing data have been made available through NCBI BioProject PRJNA807164. Individual BioSample accessions are given in Table S1. VCF files and analyses outputs are available at Dryad (will be made available upon publication). Scripts can be found at https://github.com/t-vane/ResearchSupplements (will be made available upon publication).

## Notes

### Competing Interest Statement

The authors have declared no competing interest.

