## Supplementary Text and Figures for "Species-specific responses to paleoclimatic changes and landscape barriers drive contrasting phylogeography of co-distributed lemur species in northeastern Madagascar"

### 1 Supplementary methods

#### 1.1 Restriction-site associated DNA sequencing (RADseq)

RADseq libraries were generated with the *SbfI* restriction enzyme following three different approaches (detailed for each sample in Table S4.1):

(1) Libraries were prepared according to the protocol described in Ali et al. (2016) at the University of Idaho and sequenced on an Illumina HiSeq 4000 (paired-end 150 bp) at the Vincent J. Coates Genomic Sequencing Laboratory of the University of California Berkeley, USA.

(2) Libraries were prepared according to the protocol described in Genomic Resources Development Consortium et al. (2015) and sequenced on an Illumina HiSeq 2000 (single-end, 100 bp) at the University of Oregon Core Facility, USA.

(3) Library preparation and sequencing were performed with the following custom protocol at Biosearch Technologies (LGC Genomics GmbH, Berlin, Germany). 50 ng of genomic DNA were digested with *SbfI* in 12 µl CutSmart® buffer (New England Biolabs; NEB) for 30 min at 37 °C. Restriction digests were mixed with 1.5 µl of barcoded forward *PstI* adaptors and 20 µl ligation master mix (containing 15 µl and 0.4 µl NEB Quick Ligation™ buffer and ligase, respectively). Ligation reactions were incubated for 35 min at 25°C (first ligation step). Subsequently, reactions were diluted with 30 µl Tris buffer, mixed with 50 µl Agencourt XP beads (Beckman Coulter), incubated for 15 min at room temperature, and placed for 5 min on a magnet to collect the beads. The supernatant was discarded, and the beads were washed twice with 200 µl 80% ethanol. Beads were air dried for 10 min and purified DNA was eluted in 10 µl Tris Buffer (first purification step). Subsequently, 7.5 µl of purified DNA was mixed with 7.5 µl of a master mix containing 70% Fragmentase buffer and 30% Fragmentase enzyme (Allegro® Genotyping Kit, Tecan/NuGen). Reactions were incubated for 20 min at 25°C (fragmentation/end repair step). Reaction products were again mixed with 30 µl ligation master mix (containing 22.5 µl and 0.6 µl NEB Quick Ligation™ buffer and ligase, respectively, and 10 pM standard blunt reverse adaptors). Ligation reactions were incubated for 35 min at 25°C (second ligation step). Reactions were then diluted with 20 µl Tris buffer, mixed with 50 µl Agencourt XP beads (Beckman Coulter), incubated for 15 min at room temperature, and placed for 5 min on a magnet to collect the beads. The supernatant was discarded, and the beads were washed twice with 200 µl 80% ethanol. Beads were air dried for 10 min and purified DNA was eluted in 10 µl Tris buffer (second purification step). Libraries were amplified during 19 cycles in 20 µl PCR reactions using MyTaq™ polymerase and mix (Bioline) and standard TruSeq amplification primers (Illumina). Finally, 5 µl from each of 96 amplified libraries were pooled. PCR primer and small amplicons were removed by Agencourt XP bead purification using one volume of beads. The PCR enzyme was removed by an additional purification on MinElute Columns (Qiagen). The pooled library was eluted in a final volume of 20 µl Tris buffer, size-selected on an LMP agarose gel (removing fragments smaller than 300 bp and larger than 650 bp) and sequenced on an Illumina NovaSeq 6000 (paired-end, 150 bp).

#### 1.2 Drivers of population genetic structure

The following samples were not considered for isolation-by-resistance modeling (see also Table S1):

(1) *M. lehilahytsara* individuals south of the study region to focus on landscape effects in the focal region and avoid introducing large unsampled areas, which may bias model inference.

(2) *M. simmonsi* individuals from IRS 11 and 11a (Ambodiriana and Île Ste. Marie) as the large distributional gap between these populations and those in IRSs 15–17 suggests that the associated high genetic distances are not explained by continuous landscape variables.

(3) *M. macarthurii* and *A. mooreorum* individuals as potential reproductive barriers to their sister species may bias model inference. Sample size did not allow modeling these species separately.

Climatic niche models were based on the bioclimatic variables isothermality, temperature seasonality, maximum temperature of warmest month, minimum temperature of coldest month, annual precipitation, precipitation seasonality, precipitation of wettest and driest quarter, which were previously shown to be ecologically relevant to mouse lemurs (Kamilar et al., 2016; Karger et al., 2017). All eight variables were subjected to PCA, and only the first three PCs were selected (together explaining 93.1% of the variation) to reduce multicollinearity and control for low sample sizes. The MaxEnt-algorithm of the R package 'ENMtools' v1.1.2 (Warren et al., 2021) was then used for climatic niche model estimation, and parameters were tuned independently based on lowest AIC value, using 10,000 background points. Model validation was based on the area under the receiver operating curve (AUC) and the continuous Boyce index (CBI) with a fivefold cross validation approach in the R package ENMeval v2.0.4 (Kass et al., 2021). We did not test the role of past climatic niche suitability in explaining genetic distances (e.g., as done in Fonseca et al. 2024), because paleoclimatic models for Madagascar are associated with high uncertainty (Sordyl, 2022).

### 2 Supplementary results

#### 2.1 Species distributions

*M. jonahi* was previously only known from IRSs 9 and 10 (Schüßler et al. 2020; Poelstra et al. 2021), but was now found at all sampled sites between the Voloina River in the north and the Sandratsio River in the south (IRSs 6–14) and at elevations from 40–830 m a.s.l. (Fig. S1; Table S1). Interestingly, unlike hypothesized in van Elst et al. (2025), its sister species *M. macarthurii* seems to be confined to IRS 5, as it was not found anywhere else.

The widely distributed *M. lehilahytsara* had so far been reported from several highland sites in the northeast as well as from lower elevations at Anjiahely (formerly *M. mittermeieri*; IRS 5) and Ambavala (IRS 9; see Poelstra et al. 2021; Tiley et al. 2022), where it occurs sympatrically with *M. macarthurii* and *M. jonahi*, respectively. We found *M. lehilahytsara* individuals at most sampling sites between the Masoala peninsula and the Simianona River (IRSs 2/3–11; Fig. 2B; Table S1). However, the species does not seem to occur regularly in low-elevation regions that are also inhabited by *M. jonahi* (e.g., in IRSs 6–8, 10), except for Ambavala. In contrast, *M. lehilahytsara* was found at lowland sites in IRS 2/3 and 4, where no other *Microcebus* species was observed.

*M. simmonsi* was reported from lowland forests in IRS 11 (Ambodiriana), Île Ste. Marie (formerly *M. boraha*; IRS 11a), and three sites south of the Maningory River (IRSs 16 and 17) prior to this study (Andriamasimanana et al., 2001; Rakotondravony & Rabenandrasana, 2011; Raxworthy, 1986;

Schüßler et al., 2020). Our new sampling adds one occurrence record in IRS 15 (Befotaka) to this distribution (Fig. 2C; Table S1). Notably, we did not find any *M. simmonsi* individuals between the Simianona and Sandratsio Rivers despite sampling all three interjacent IRSs (12–14), which contained *M. jonahi* instead. This suggests a distributional gap between the northern and southern sampling sites of *M. simmonsi*.

Finally, the Antainambalana River presents a distributional boundary towards the Masoala peninsula for *A. laniger* (Figs. 2D, S2; Table S1), which has been assumed to occur along Madagascar's east coast between the Mangoro River in the south and the Bemarivo River in the north (Mittermeier et al., 2023). Mitochondrial data by Lei et al. (2008) indicate that the distribution of *A. laniger* extends north up to Anjanaharibe-Sud Special Reserve, but our sampling did not allow testing this hypothesis (Fig. S2). We show, however, that IRS 4 is occupied by *A. mooreorum* instead (Figs. 2D, S2; Table S1), previously only known around its type locality in IRS 2/3 (Lei et al., 2008; Mittermeier et al., 2023).

### 3 Supplementary figures

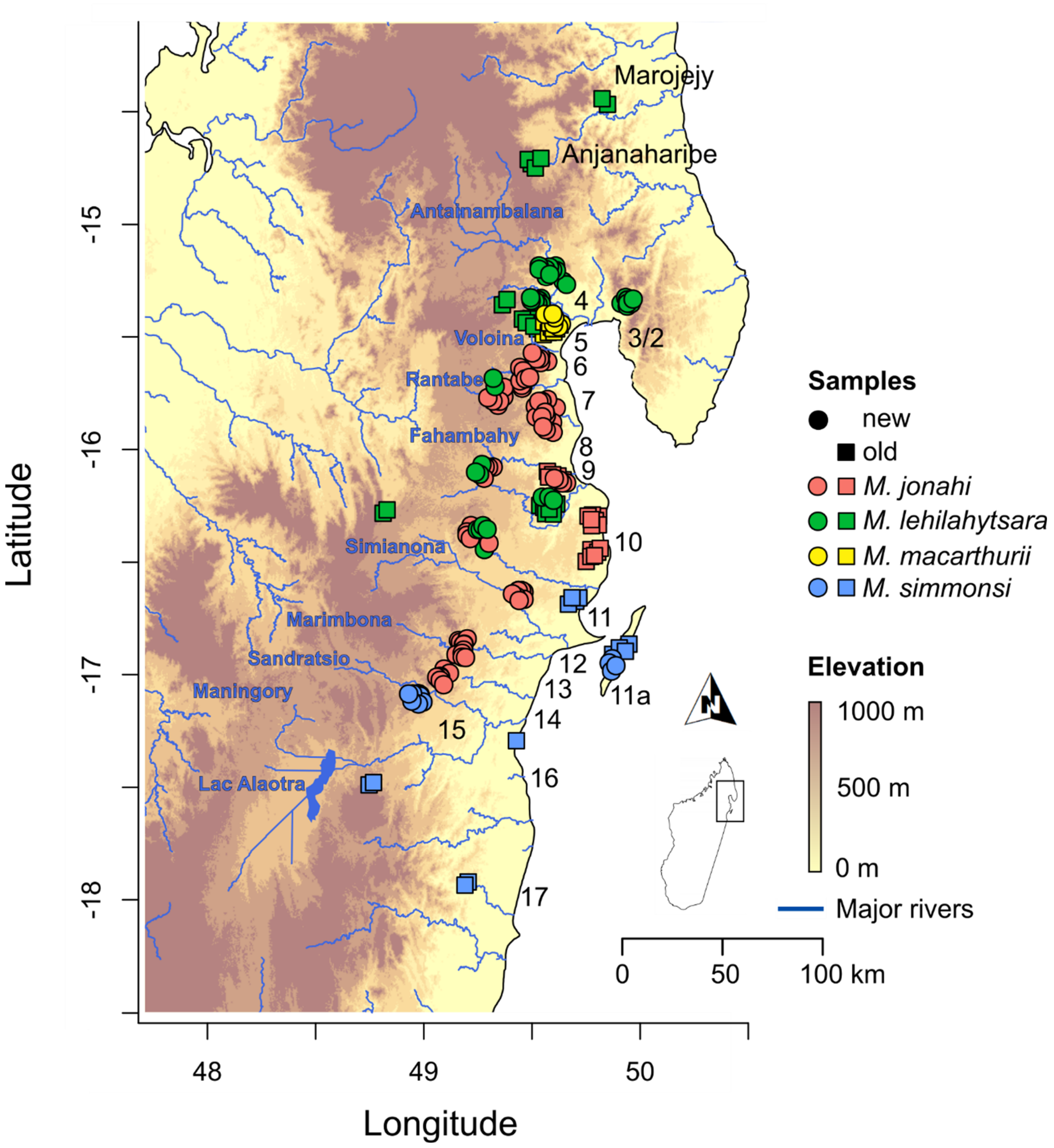


**Fig. S1:** Sampled *Microcebus* spp. individuals across the study region in northeastern Madagascar. Sample information is given in Table S1. Numbers denote inter-river systems. Blue labels denote names of rivers and Lac Alaotra.

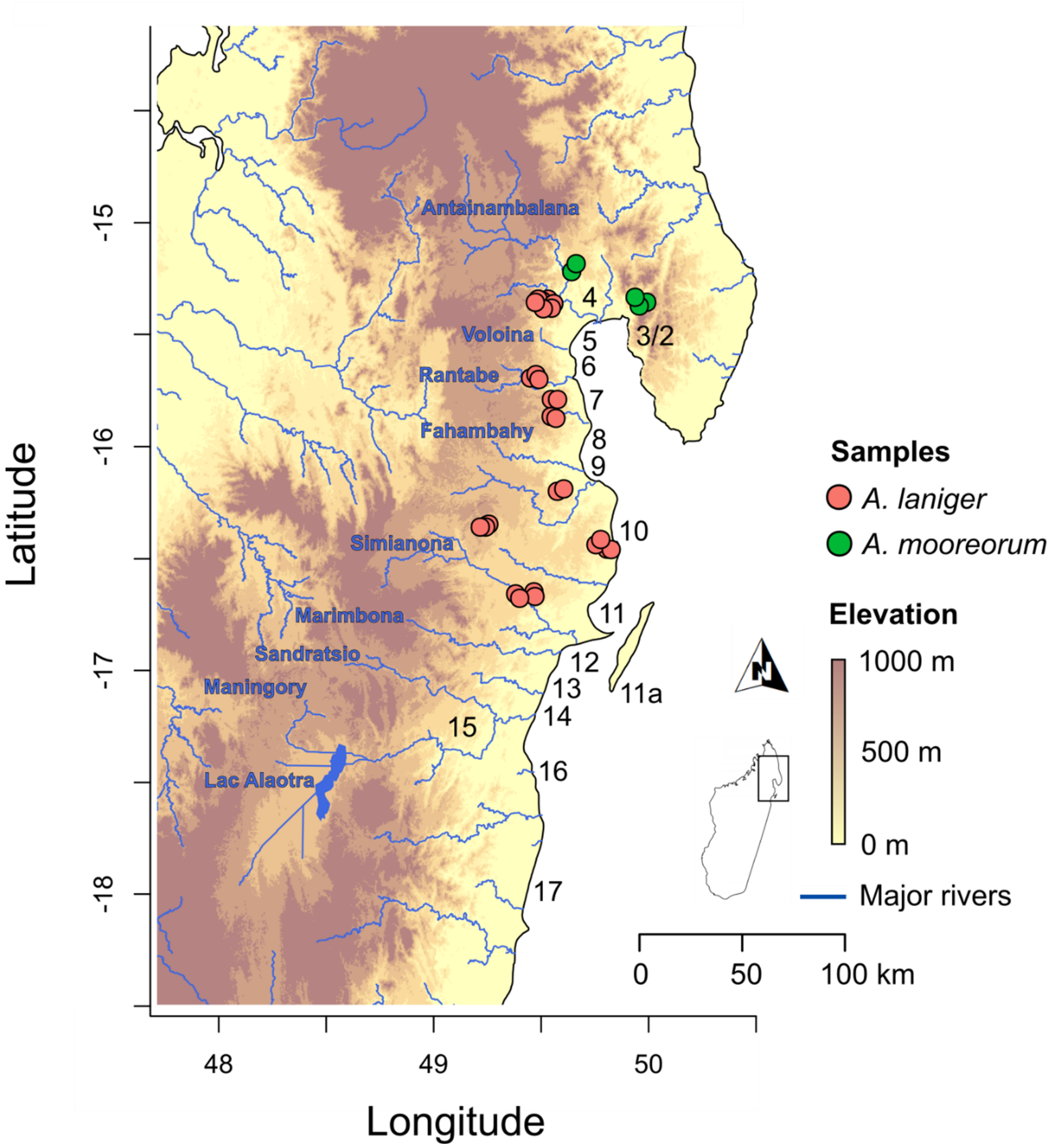


**Fig. S2:** Sampled *Avahi* spp. individuals across the study region in northeastern Madagascar. Sample information is given in Table S1. Numbers denote inter-river systems. Blue labels denote names of rivers and Lac Alaotra.

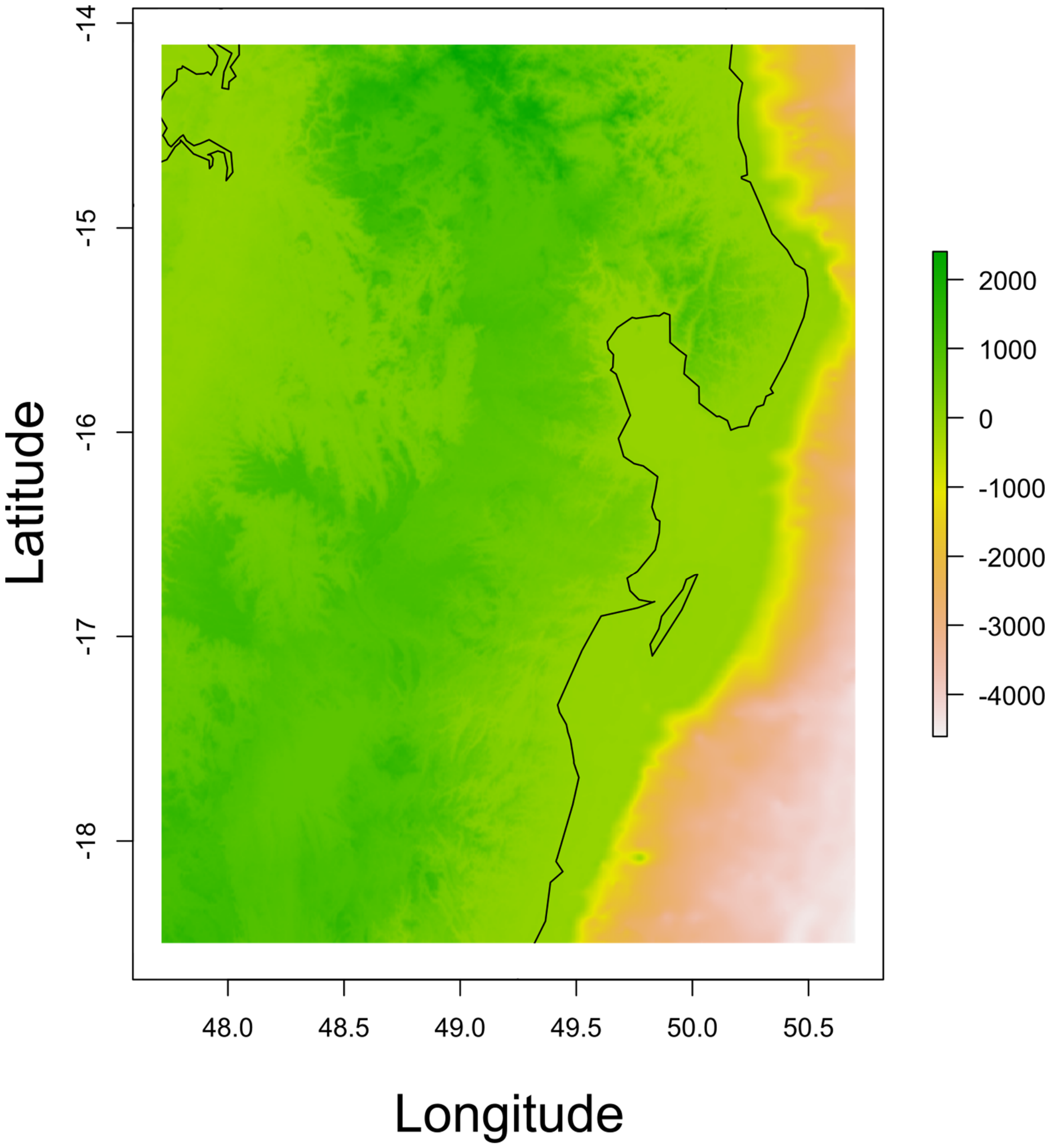


**Fig. S3:** Elevation (m) raster across the entire study region (not scaled) used for isolation-by-resistance analysis. Resolution is 150 m per pixel.

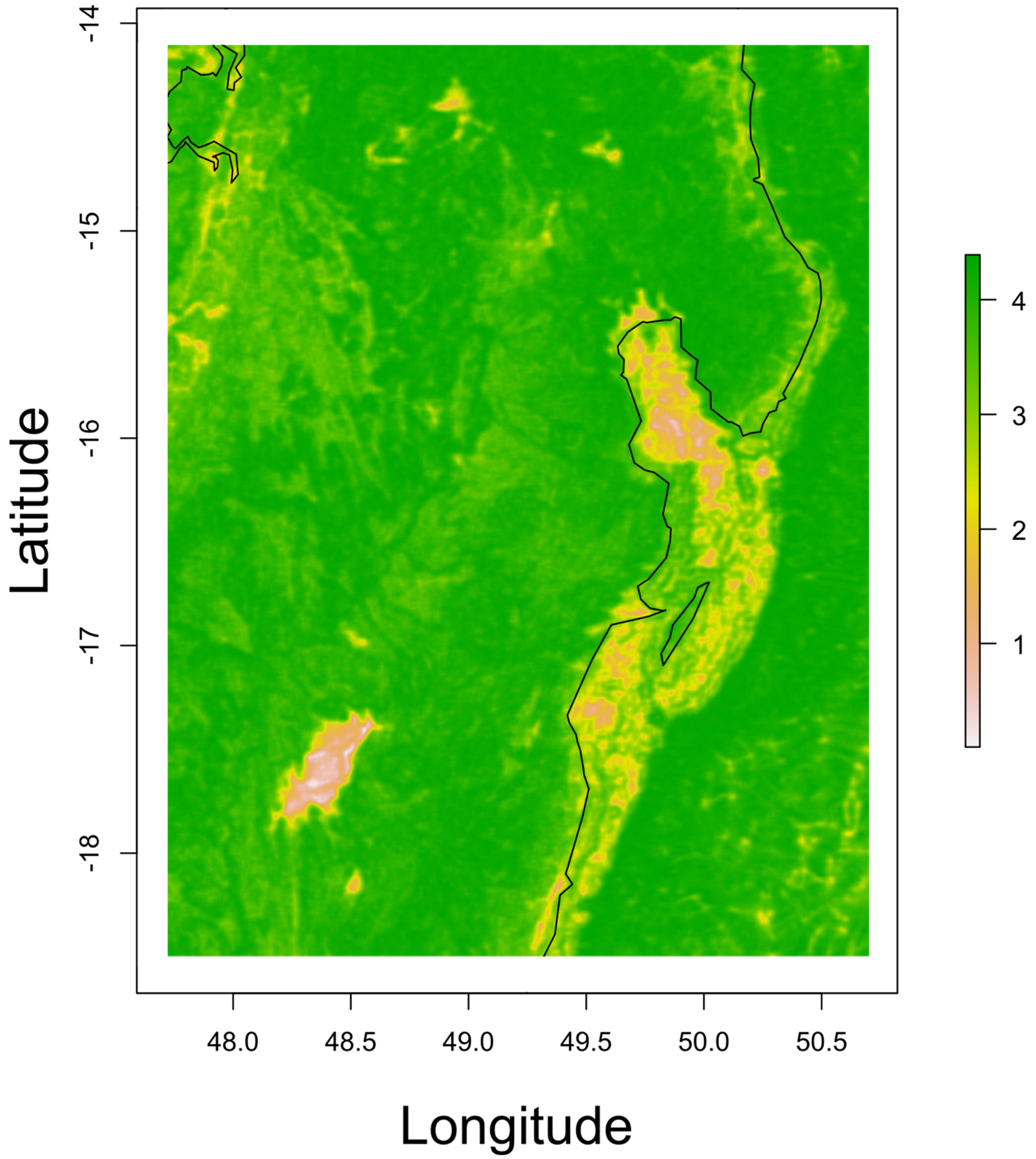


**Fig. S4:** Landscape heterogeneity raster across the entire study region (not scaled) used for isolation-by-resistance analysis. Resolution is 150 m per pixel.

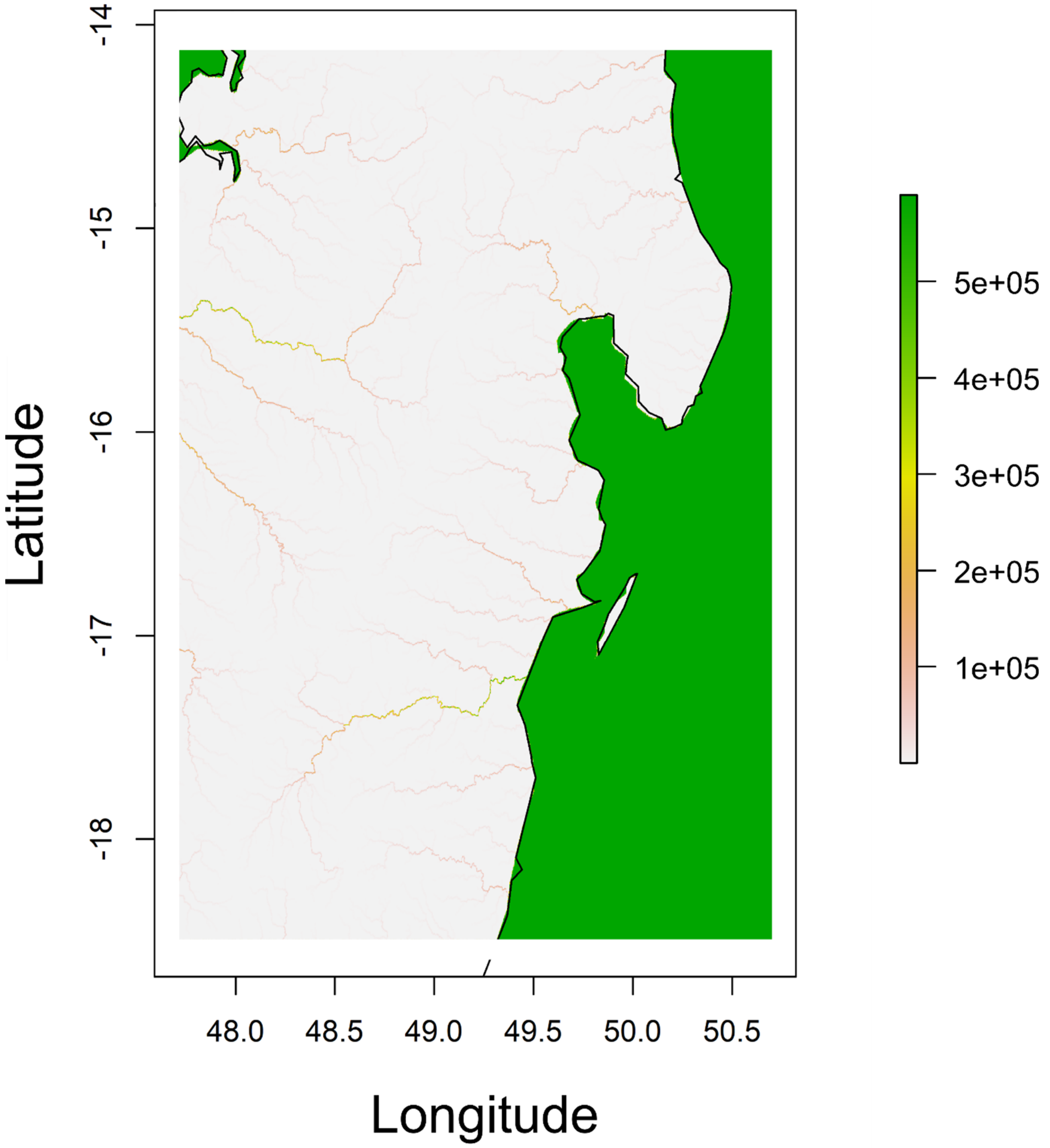


**Fig. S5:** Flow accumulation raster across the entire study region (current sea level; not scaled) used for isolation-by-resistance analysis. Resolution is 150 m per pixel. River width has increased for visibility.

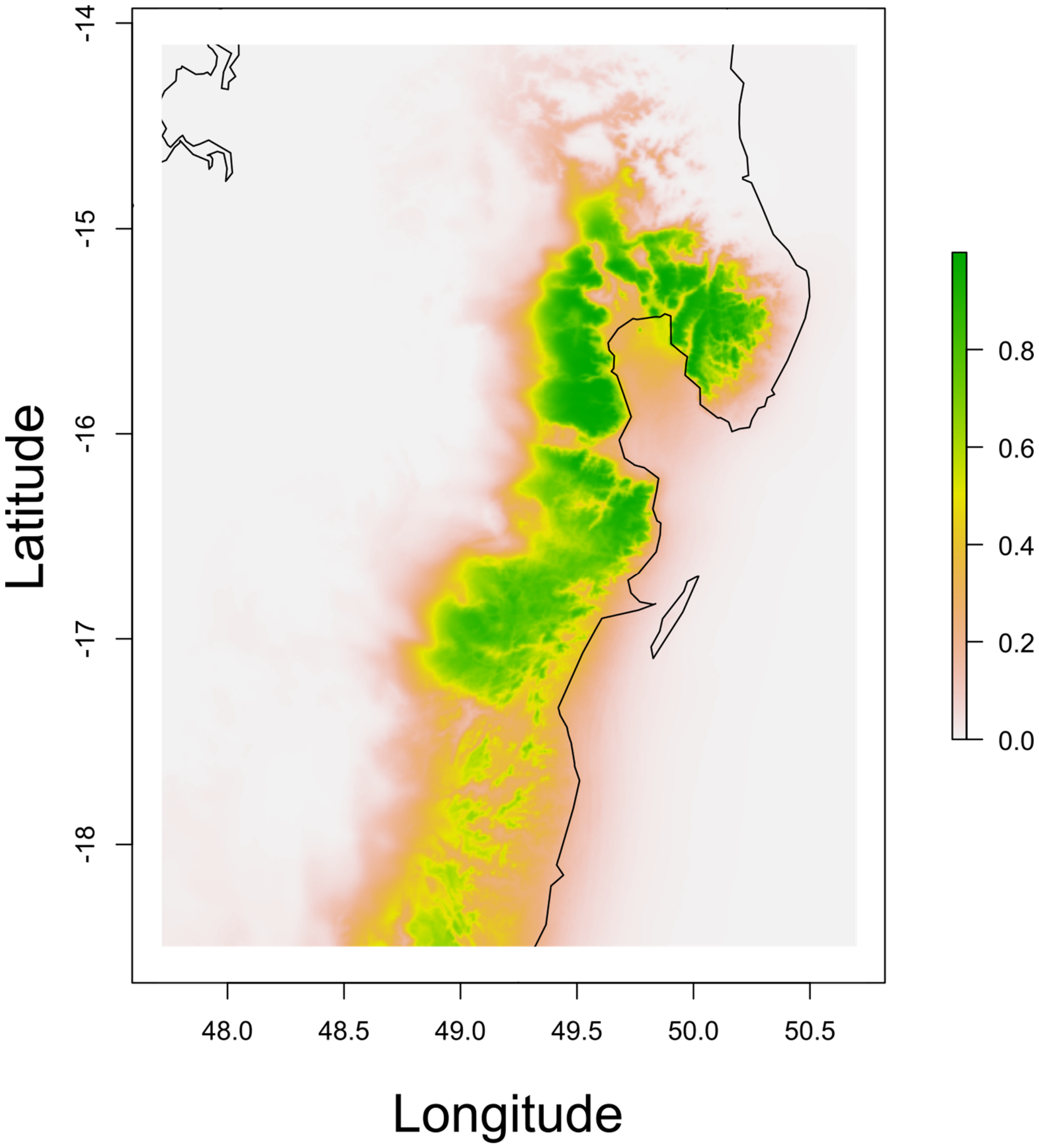


**Fig. S6:** Climatic niche suitability raster for *M. jonahi* across the entire study region (not scaled) used for isolation-by-resistance analysis. Resolution is 150 m per pixel.

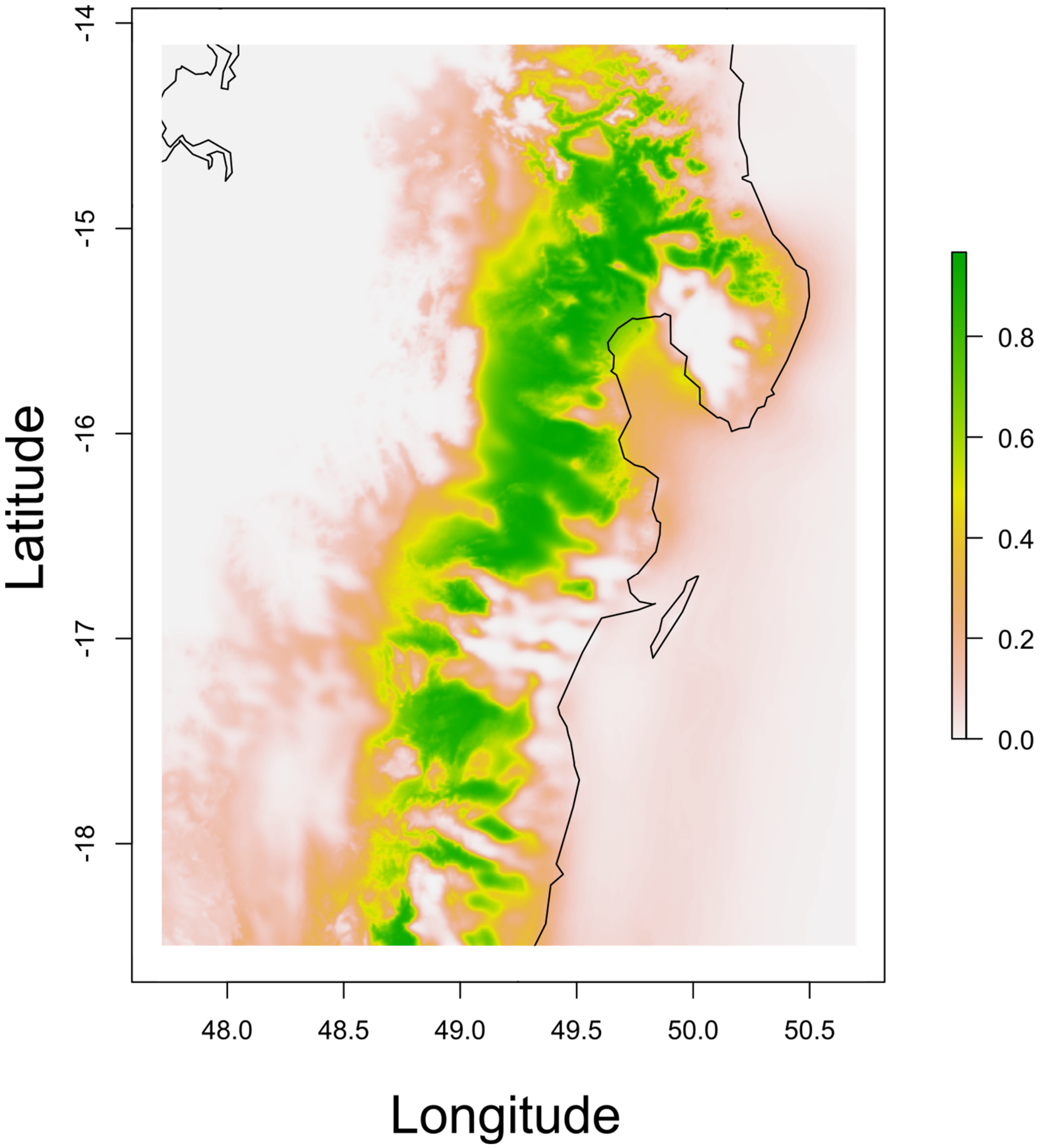


**Fig. S7:** Climatic niche suitability raster for *M. lehilahytsara* across the entire study region (not scaled) used for isolation-by-resistance analysis. Resolution is 150 m per pixel.

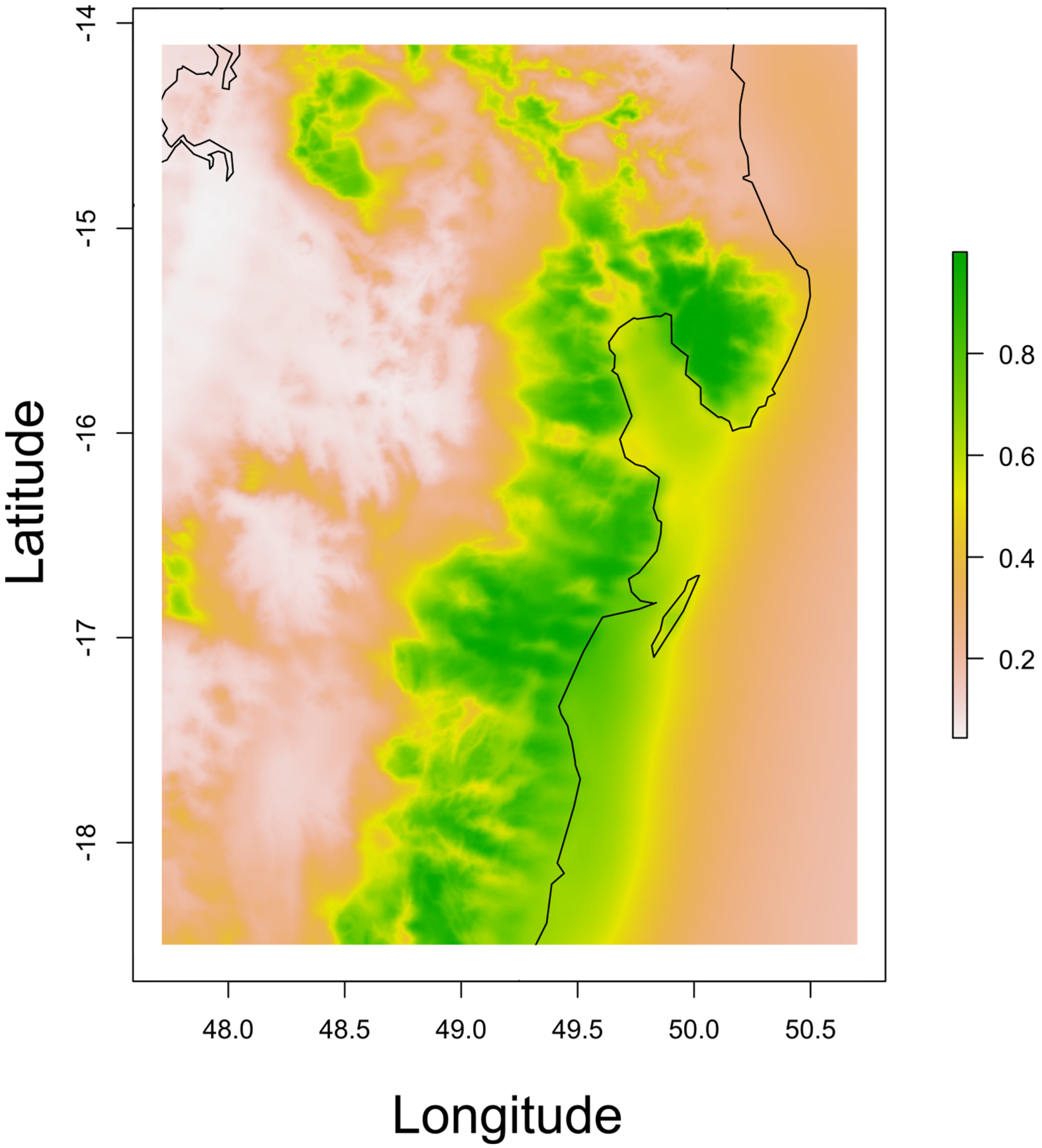


**Fig. S8:** Climatic niche suitability raster for *M. simmonsi* across the entire study region (not scaled) used for isolation-by-resistance analysis. Resolution is 150 m per pixel.

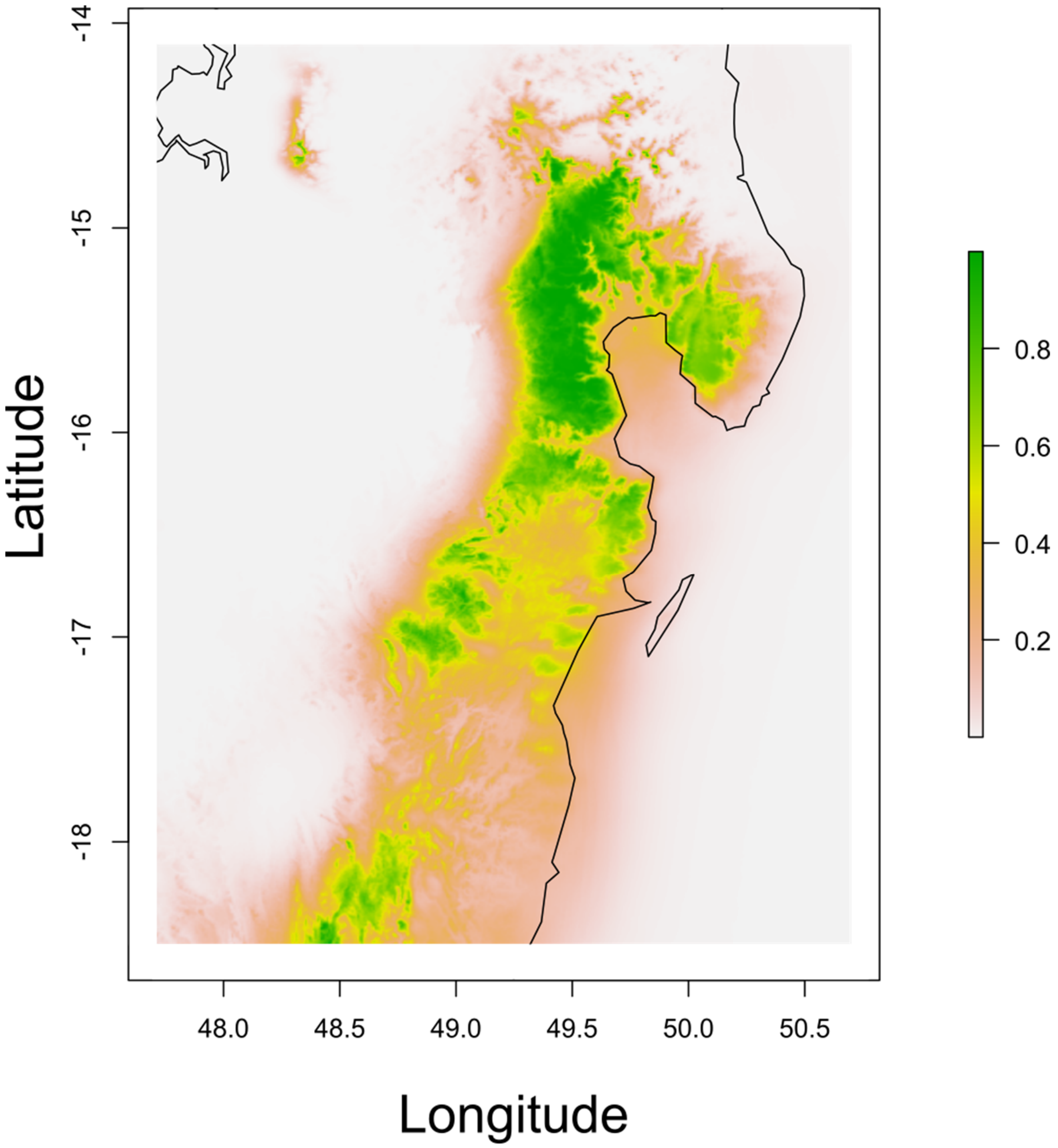


**Fig. S9:** Climatic niche suitability raster for *A. laniger* across the entire study region (not scaled) used for isolation-by-resistance analysis. Resolution is 150 m per pixel.

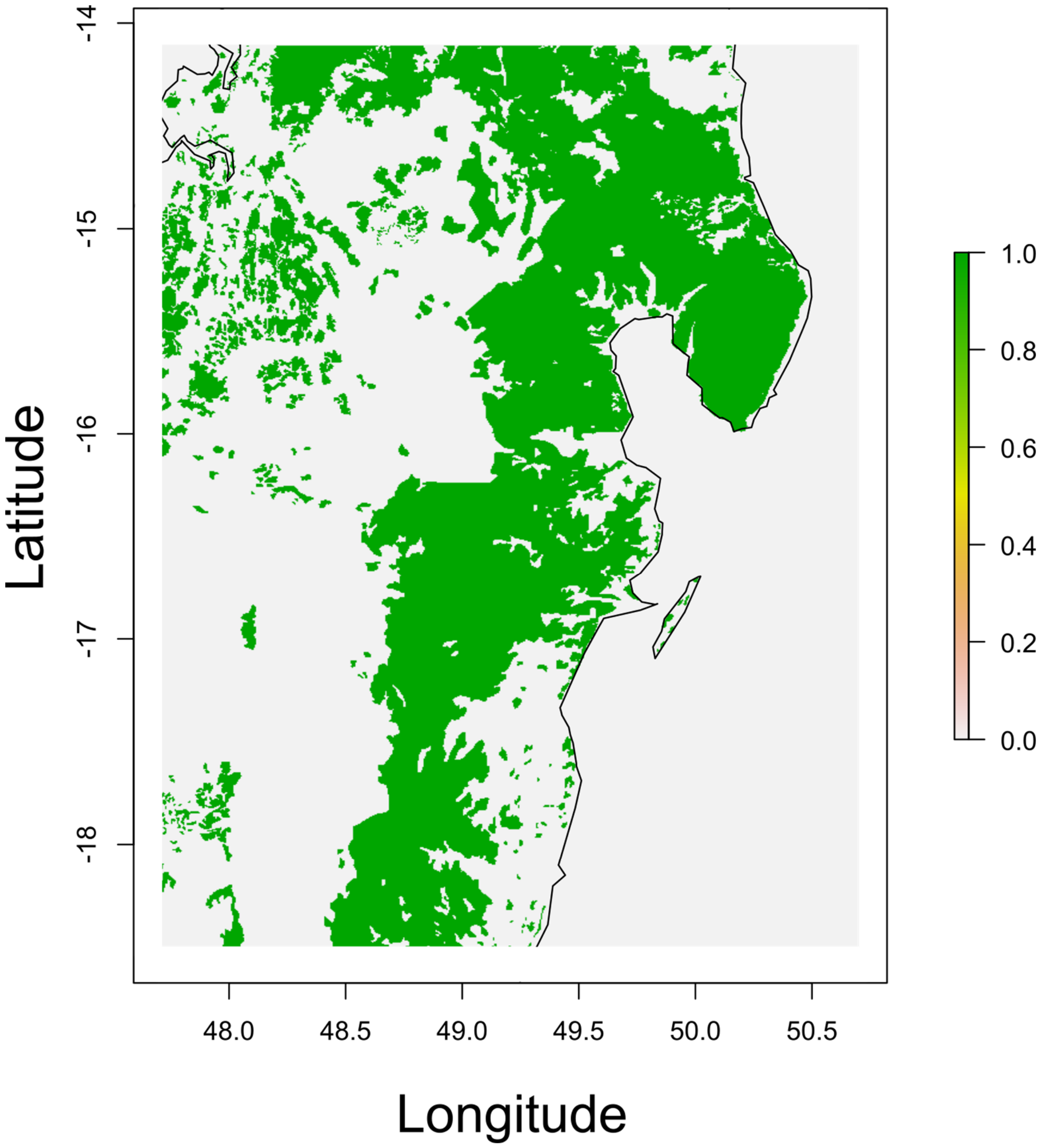


**Fig. S10:** Raster corresponding to forest cover at 1953 (Vieilledent et al., 2018) across the entire study region used for isolation-by-resistance analysis. Resolution is 150 m per pixel.

*M. jonahi*

| | Elevation | Landscape heterogeneity | Rivers | Climatic niche suitability | Forest |
|---|---|---|---|---|---|
| Elevation | 1.00 | 0.30 | -0.40 | -0.40 | 0.57 |
| Landscape heterogeneity | 0.30 | 1.00 | -0.23 | 0.28 | -0.07 |
| Rivers | -0.40 | -0.23 | 1.00 | -0.13 | -0.01 |
| Climatic niche suitability | -0.40 | 0.28 | -0.13 | 1.00 | 0.57 |
| Forest | 0.57 | -0.07 | -0.01 | 0.57 | 1.00 |

*M. lehilahytsara*

| | Elevation | Landscape heterogeneity | Rivers | Climatic niche suitability | Forest |
|---|---|---|---|---|---|
| Elevation | 1.00 | 0.32 | -0.27 | -0.04 | 0.13 |
| Landscape heterogeneity | 0.32 | 1.00 | -0.12 | -0.16 | 0.26 |
| Rivers | -0.27 | -0.12 | 1.00 | -0.02 | -0.10 |
| Climatic niche suitability | -0.04 | -0.16 | -0.02 | 1.00 | 0.47 |
| Forest | 0.13 | 0.26 | -0.10 | 0.47 | 1.00 |

*M. simmonsi*

| | Elevation | Landscape heterogeneity | Rivers | Climatic niche suitability | Forest |
|---|---|---|---|---|---|
| Elevation | 1.00 | -0.13 | -0.66 | -0.36 | 0.47 |
| Landscape heterogeneity | -0.13 | 1.00 | -0.17 | -0.11 | 0.22 |
| Rivers | -0.66 | -0.17 | 1.00 | -0.27 | 0.03 |
| Climatic niche suitability | -0.36 | -0.11 | -0.27 | 1.00 | 0.39 |
| Forest | 0.47 | 0.22 | 0.03 | 0.39 | 1.00 |

*A. laniger*

| | Elevation | Landscape heterogeneity | Rivers | Climatic niche suitability | Forest |
|---|---|---|---|---|---|
| Elevation | 1.00 | 0.18 | -0.37 | -0.14 | 0.34 |
| Landscape heterogeneity | 0.18 | 1.00 | -0.22 | 0.31 | 0.09 |
| Rivers | -0.37 | -0.22 | 1.00 | -0.04 | -0.08 |
| Climatic niche suitability | -0.14 | 0.31 | -0.04 | 1.00 | 0.53 |
| Forest | 0.34 | 0.09 | -0.08 | 0.53 | 1.00 |

**Fig. S11:** Correlation coefficients for pairwise comparisons of cropped and scaled rasters at a resolution of 150 m for each species (based on a pixel subsample of 5%). Red and blue colors indicate negative and positive correlations, respectively. Color intensity is proportional to the strength of the correlation.

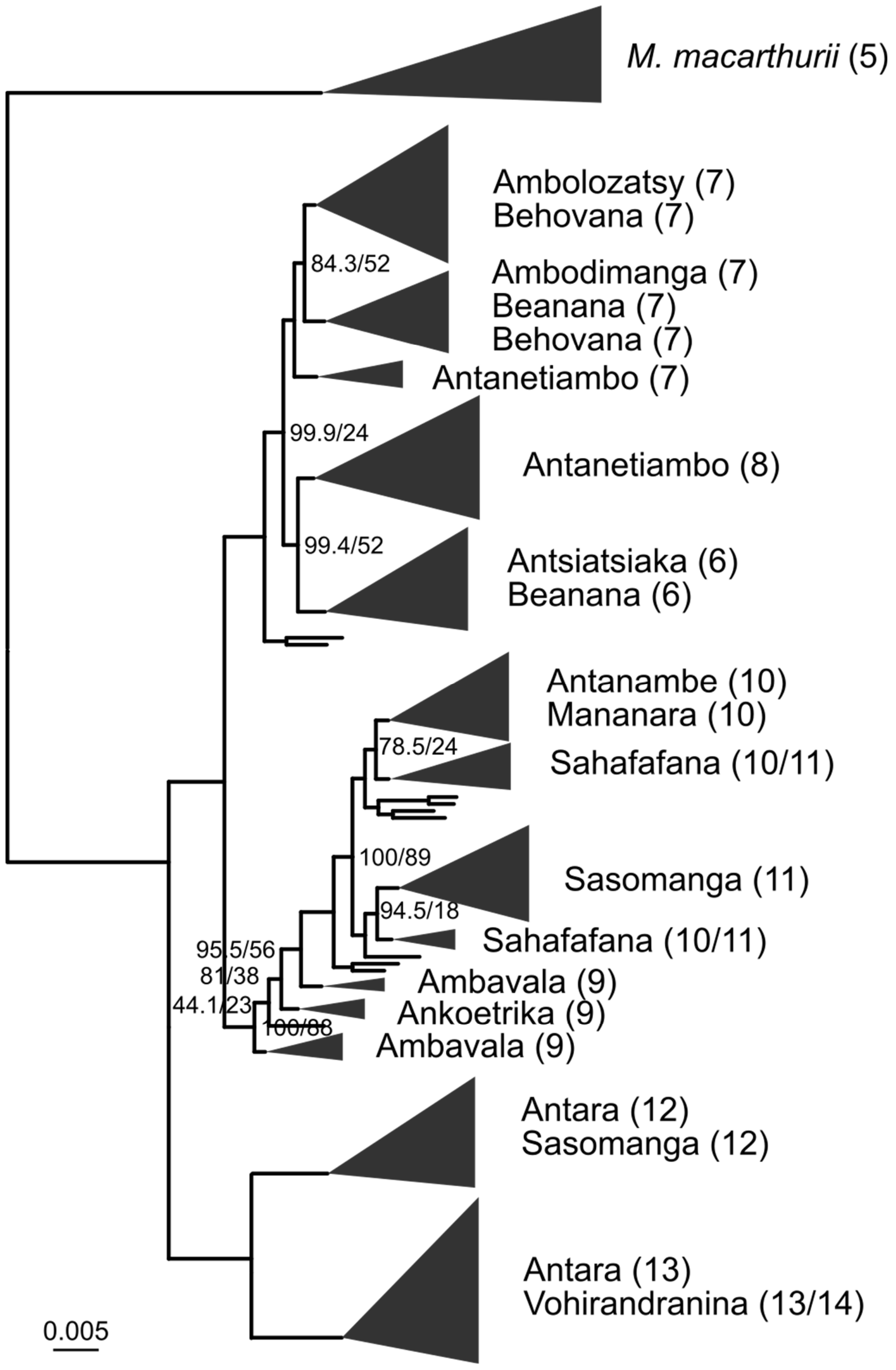


**Fig. S12:** Maximum likelihood phylogeny inferred with IQ-TREE for *M. jonahi* and *M. macarthurii*. Tip labels denote population names (of *M. jonahi* if not mentioned otherwise) and associated inter-river systems (in parentheses). Triangles represent collapsed tips proportional to sample size. Node labels represent ultrafast bootstrap and SH-like approximate likelihood ratio (SH-aLRT) support, given only for major clades and if below 100. Scale is substitutions per site.

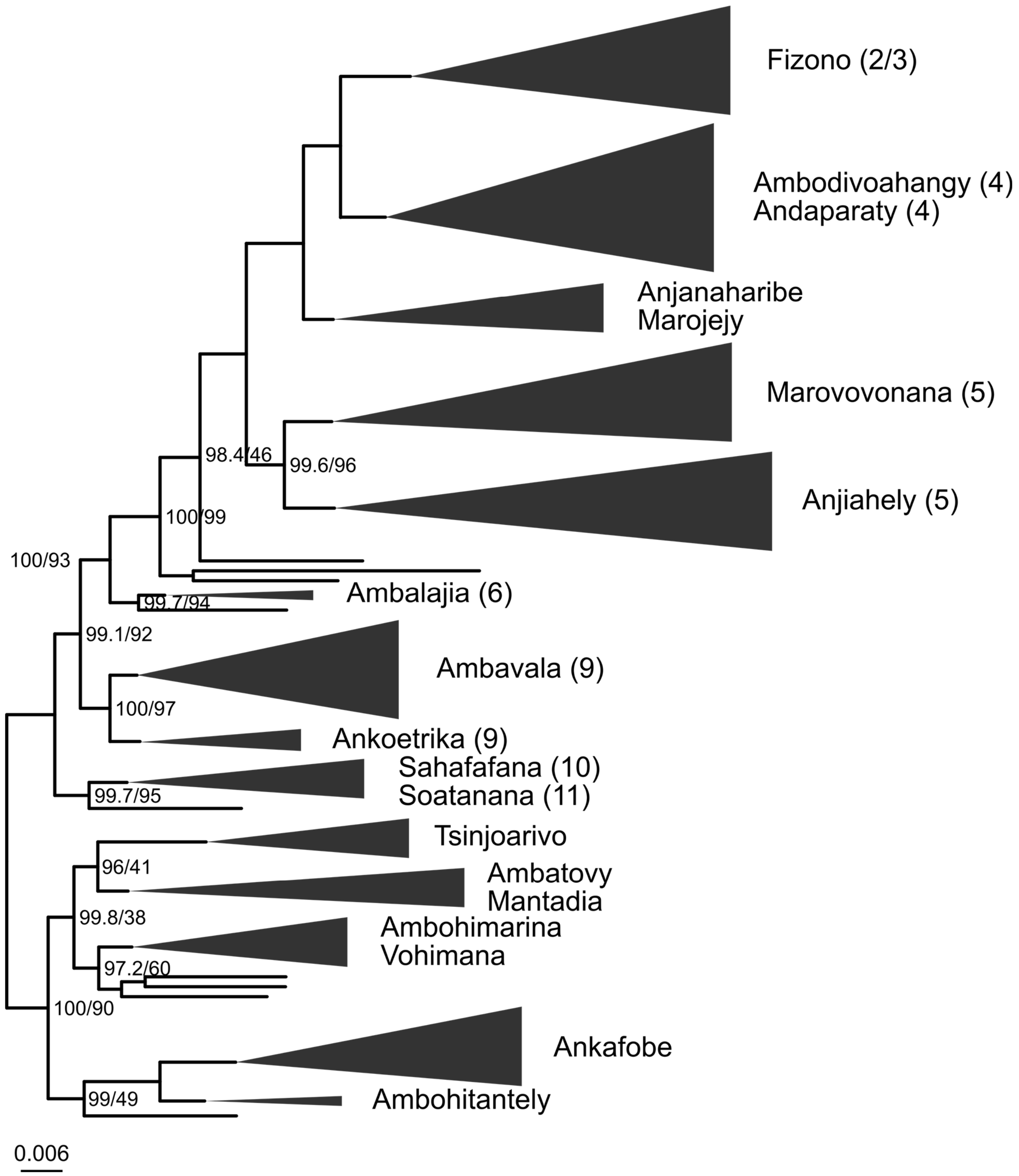


**Fig. S13:** Maximum likelihood phylogeny inferred with IQ-TREE for *M. lehilahytsara*. Tip labels denote population names and associated inter-river systems if applicable (in parentheses). Triangles represent collapsed tips proportional to sample size. Node labels represent ultrafast bootstrap and SH-like approximate likelihood ratio (SH-aLRT) support, given only for major clades and if below 100. Scale is substitutions per site.

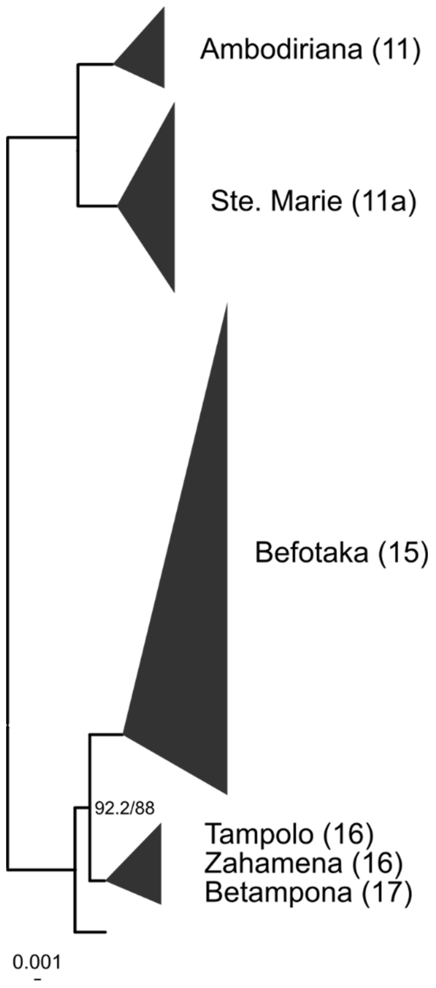


**Fig. S14:** Maximum likelihood phylogeny inferred with IQ-TREE for *M. simmonsi*. Tip labels denote population names and associated inter-river systems (in parentheses). Triangles represent collapsed tips proportional to sample size. Node labels represent ultrafast bootstrap and SH-like approximate likelihood ratio (SH-aLRT) support, given only for major clades and if below 100. Scale is substitutions per site.

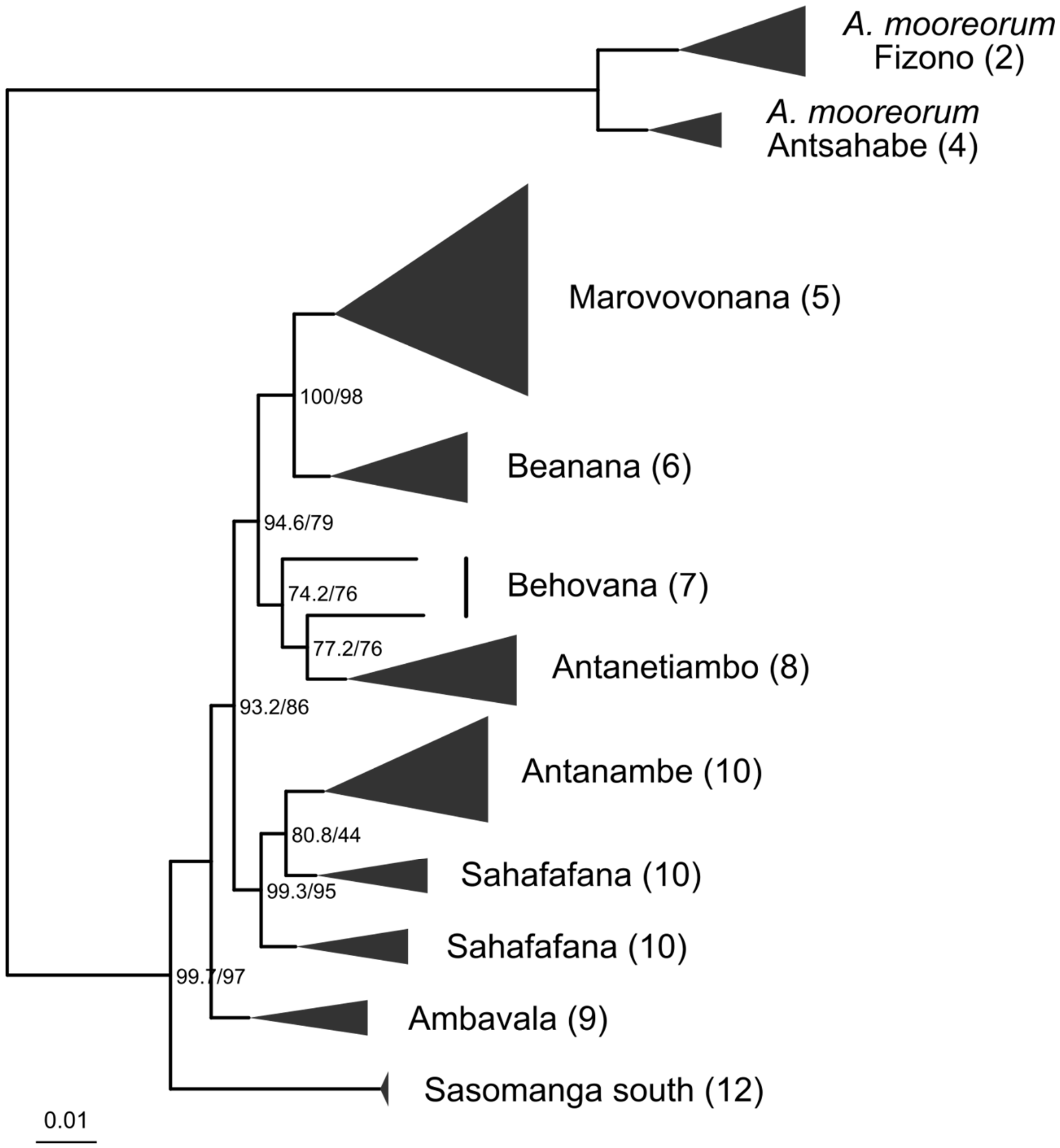


**Fig. S15:** Maximum likelihood phylogeny inferred with IQ-TREE for *A. laniger* and *A. mooreorum*. Tip labels denote population names (of *A. laniger* if not mentioned otherwise) and associated inter-river systems (in parentheses). Triangles represent collapsed tips proportional to sample size. Node labels represent ultrafast bootstrap and SH-like approximate likelihood ratio (SH-aLRT) support, given only for major clades and if below 100. Scale is substitutions per site.

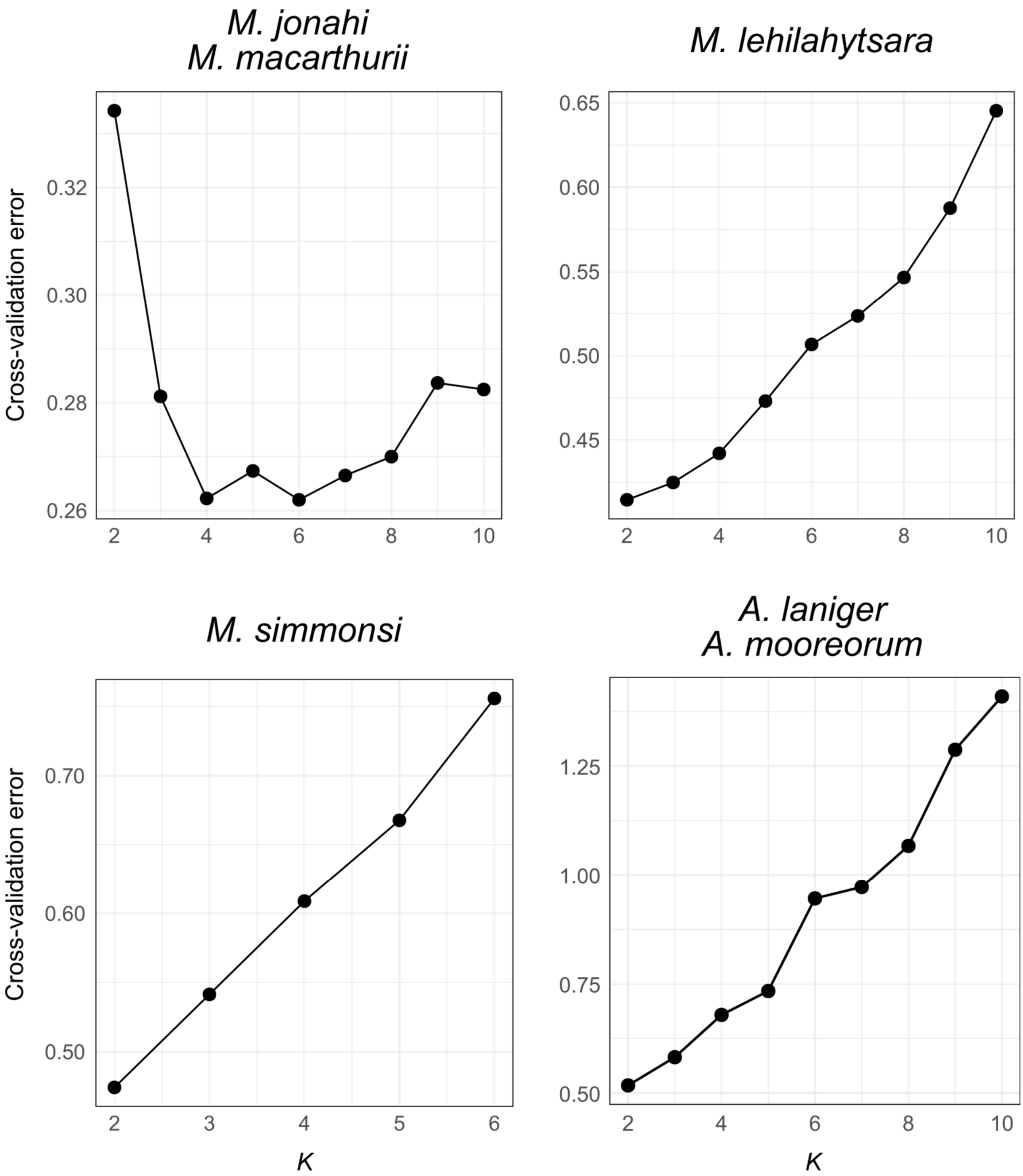


**Fig. S16:** Cross-validation errors for different numbers of clusters (*K*) in ancestry inference with ADMIXTURE.

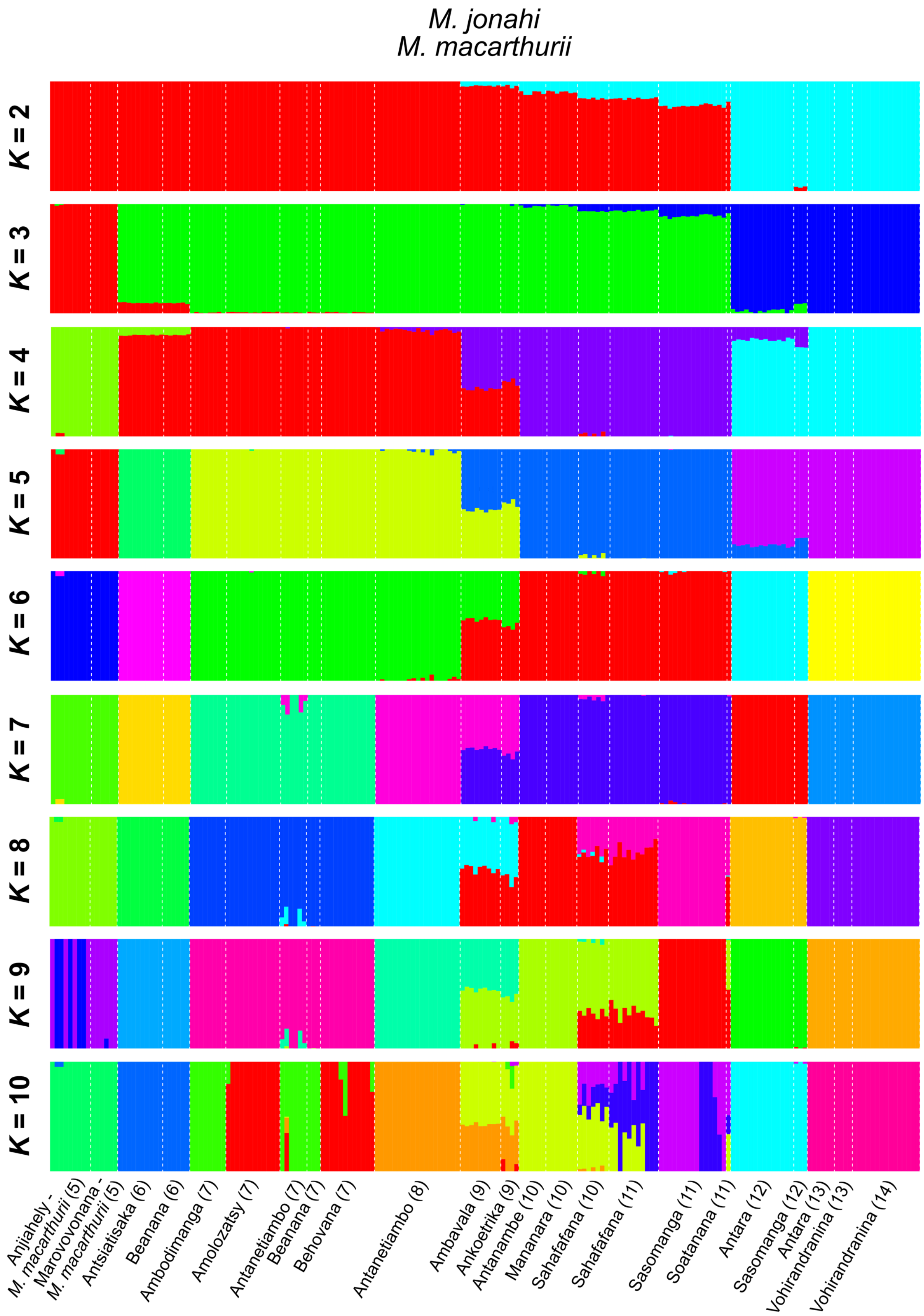


**Fig. S17:** Admixture proportions of *M. jonahi* and *M. macarthurii* individuals (columns) estimated with ADMIXTURE for two to ten clusters (*K*). Population names refer to *M. jonahi* if not mentioned otherwise. Numbers denote inter-river systems.

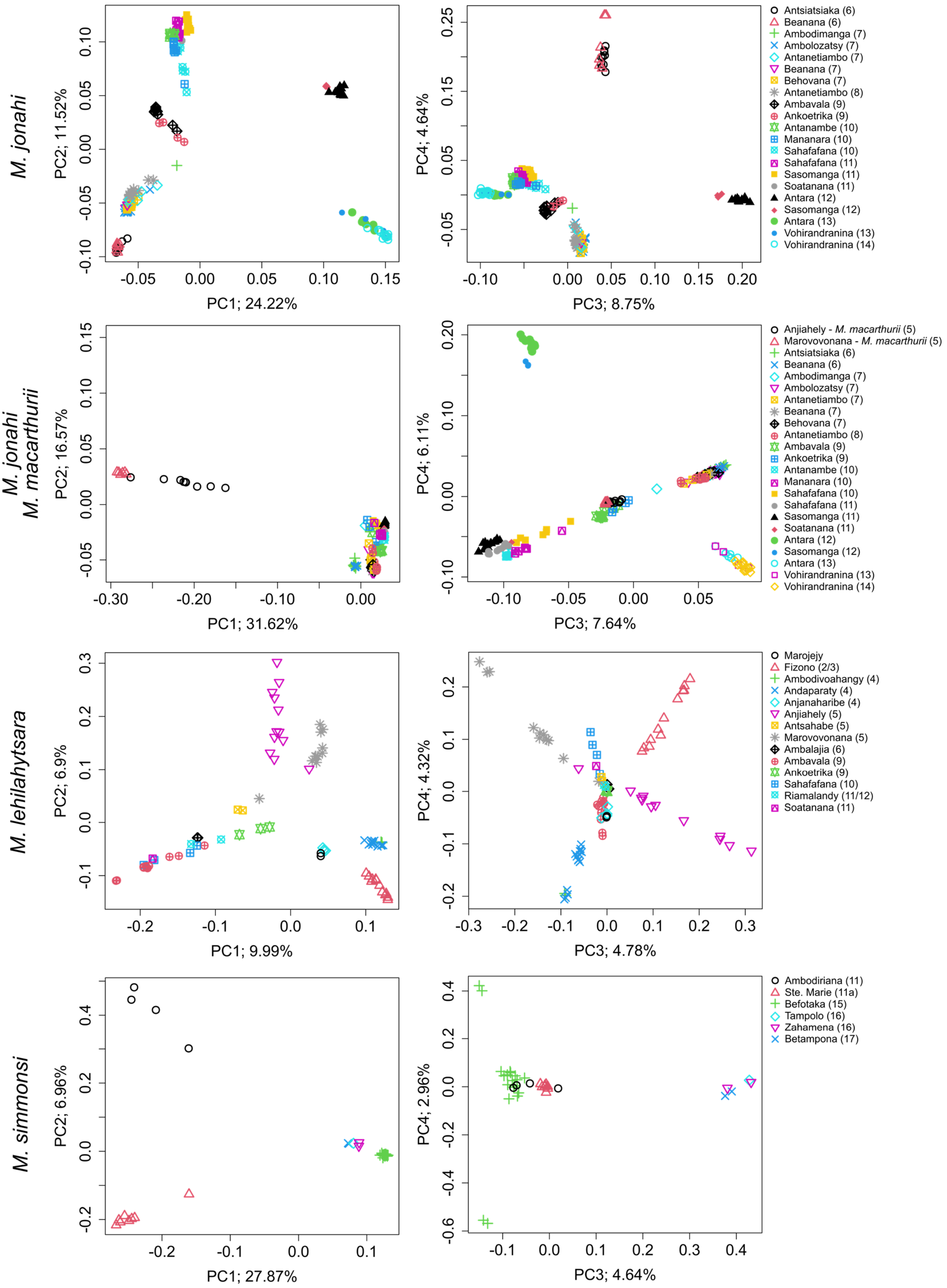


**Fig. S18:** Principal component analyses (PCA) of genetic data. Left column: PC1 plotted against PC2; Right column: PC3 plotted against PC4. Percentages indicate the amount of variation explained by the PC. In plots including *M. jonahi*, population names refer to *M. jonahi* if not mentioned otherwise. Numbers after denote inter-river systems.

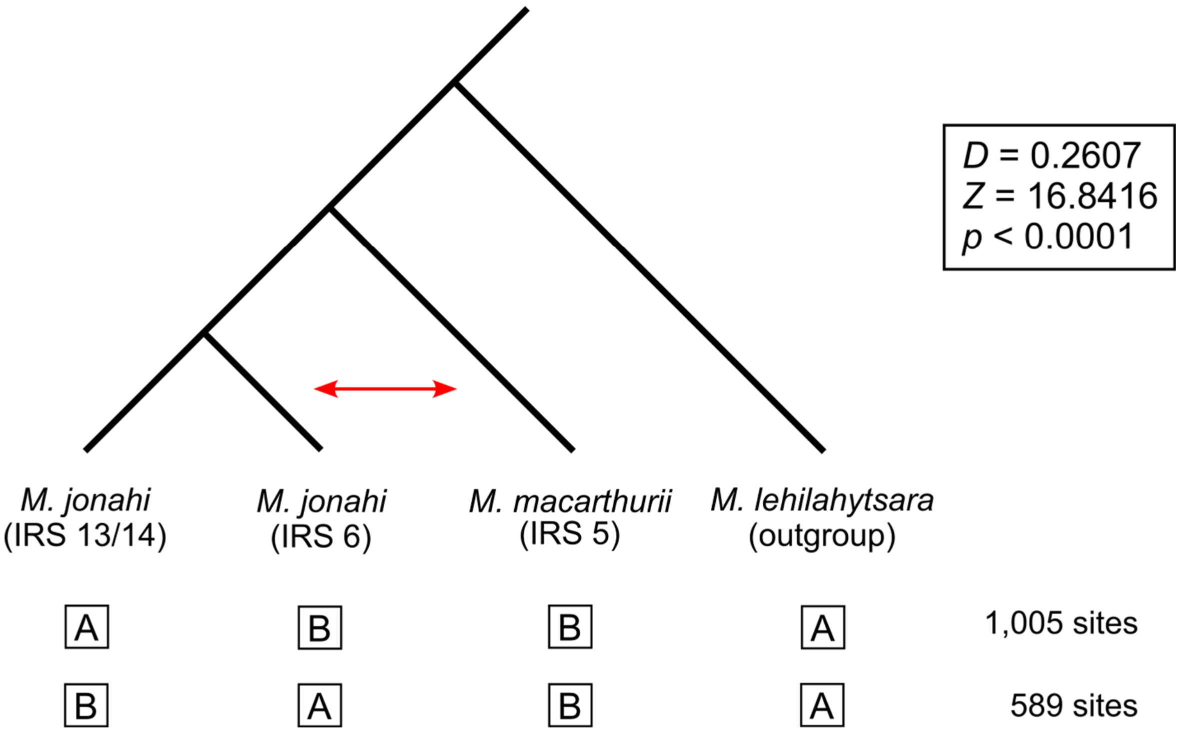


**Fig. S19:** Test for introgression between *M. macarthurii* (inter-river system/IRS 5) and the *M. jonahi* populations from IRS 6 (indicated by red arrow) via Patterson's *D* statistic. Significantly more shared sites were found between these clades (ABBA) than between *M. macarthurii* and the *M. jonahi* populations from IRSs 12 and 13 (BABA), providing strong evidence for introgression.

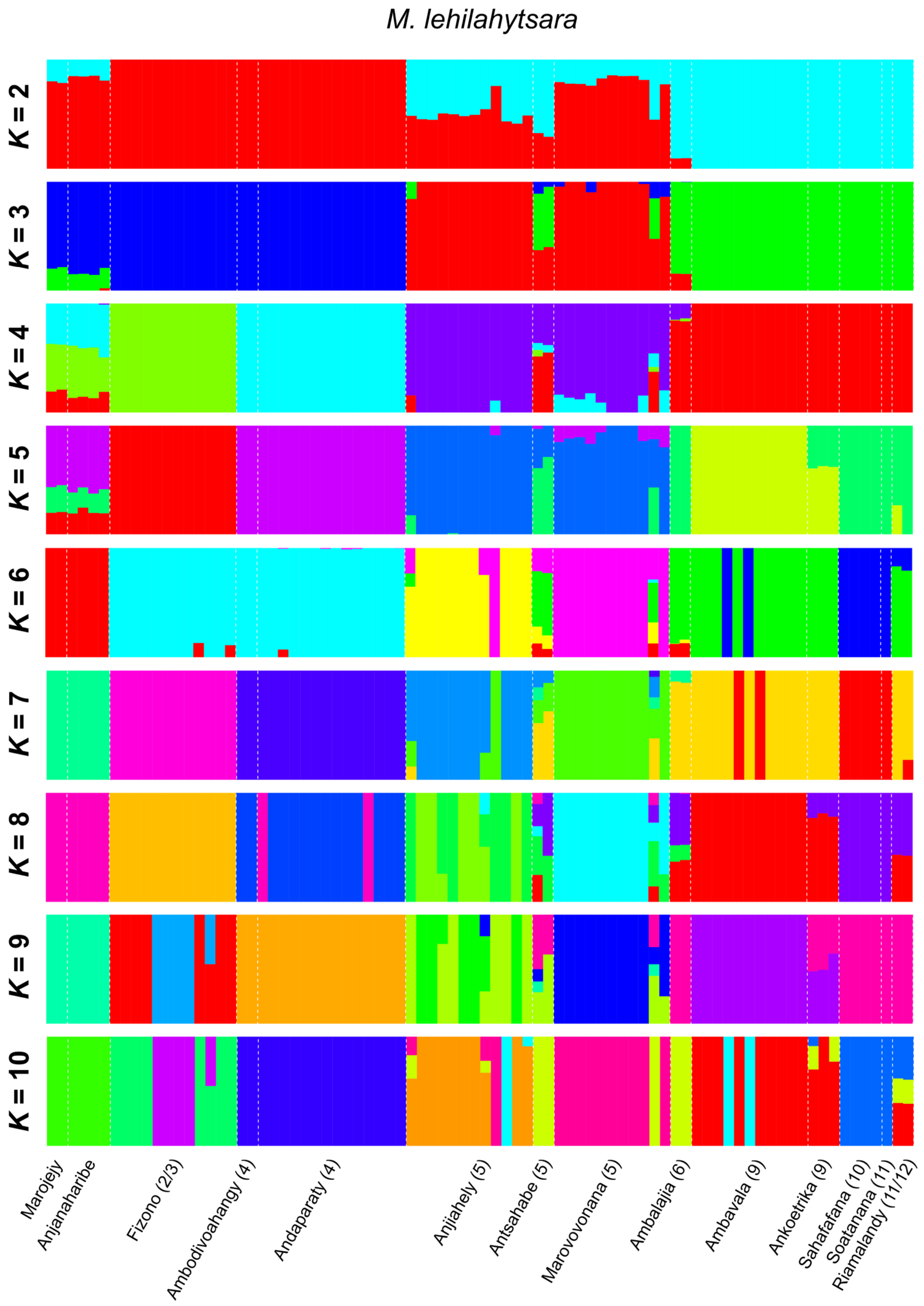


**Fig. S20:** Admixture proportions of *M. lehilahytsara* individuals (columns) estimated with ADMIXTURE for two to ten clusters (*K*). Numbers denote inter-river systems.

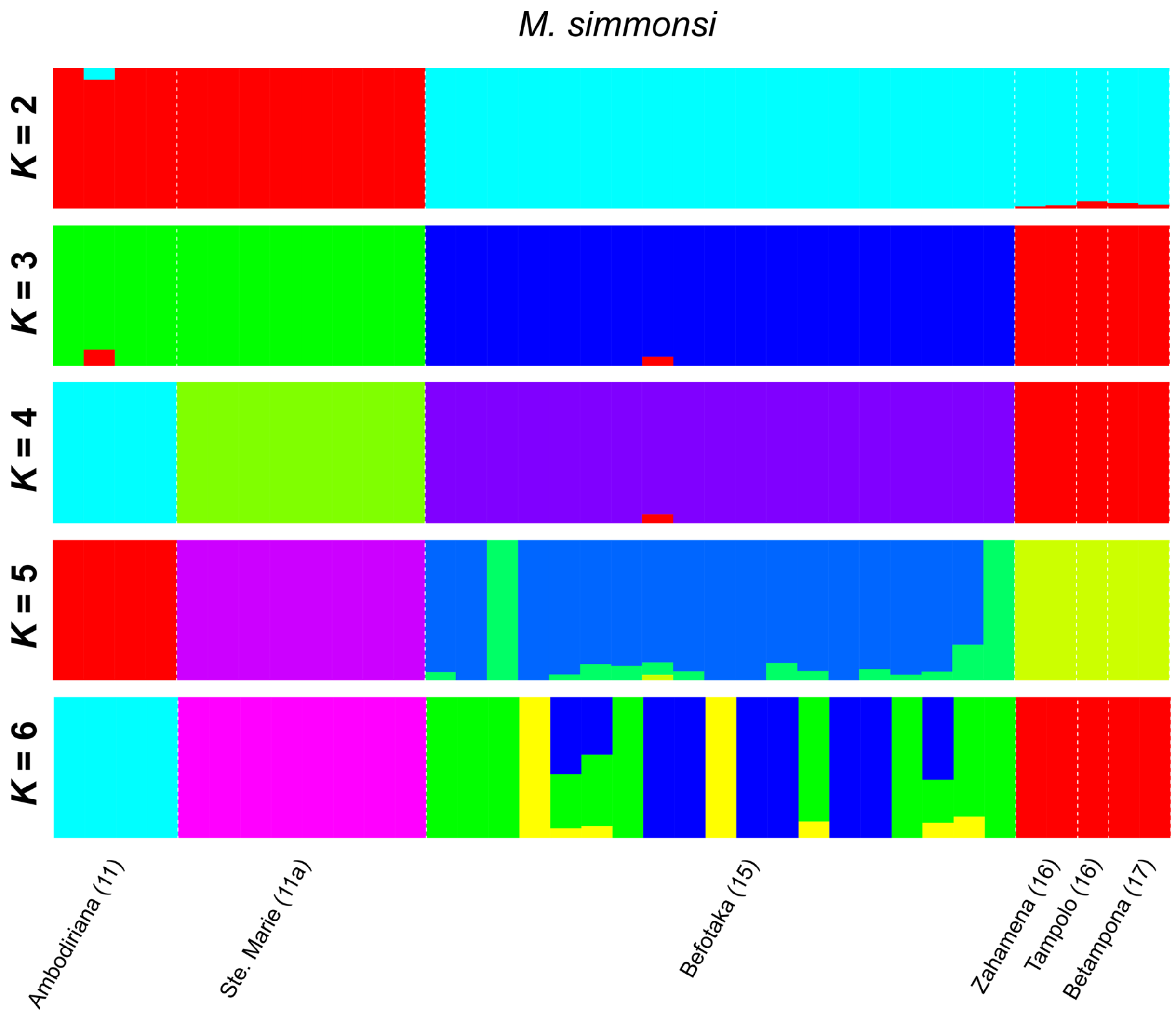


**Fig. S21:** Admixture proportions of *M. simmonsi* individuals (columns) estimated with ADMIXTURE for two to six clusters (*K*). Numbers denote inter-river systems.

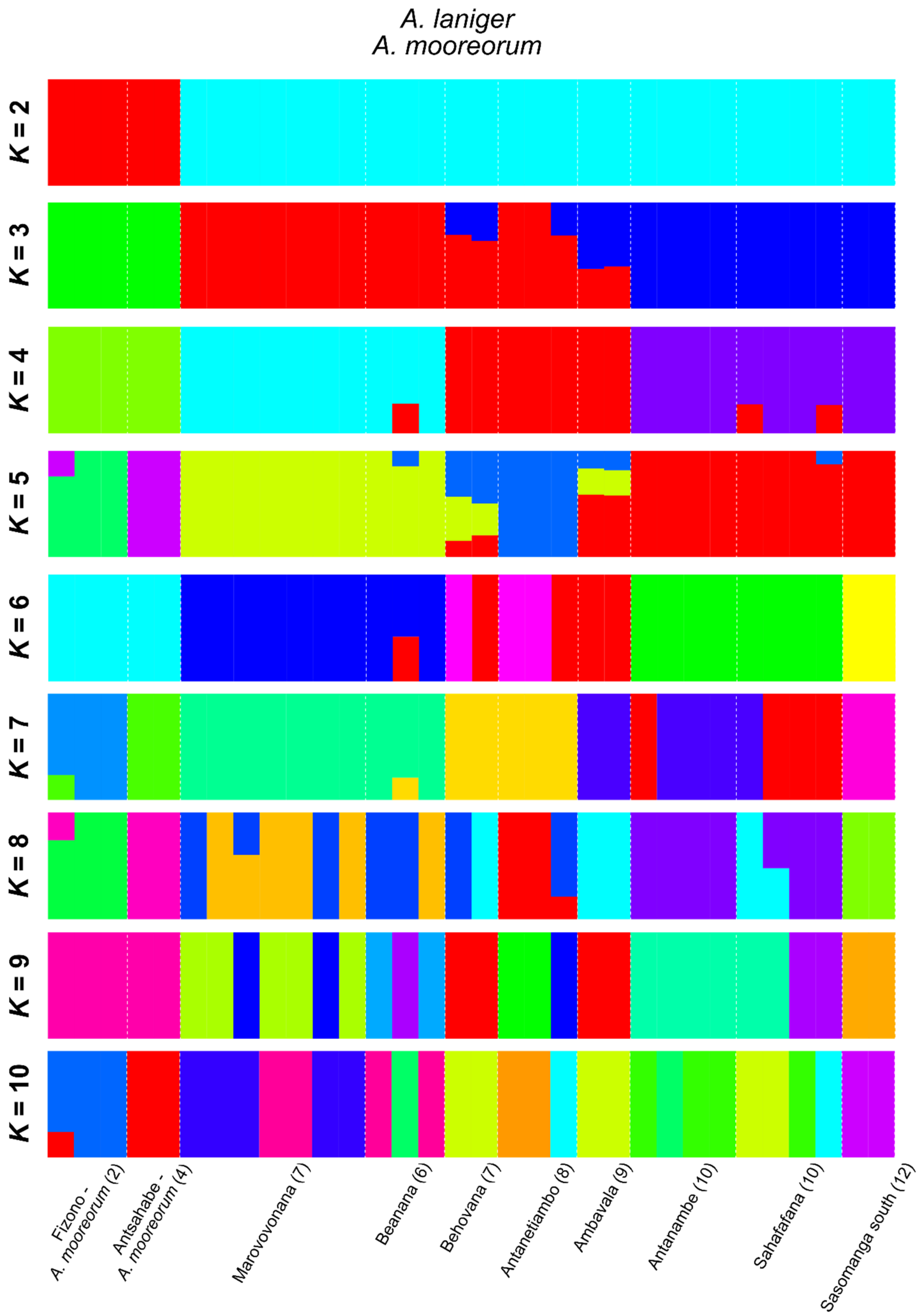


**Fig. S22:** Admixture proportions of *A. laniger* and *A. mooreorum* individuals (columns) estimated with ADMIXTURE for two to ten clusters (*K*). Population names refer to *A.laniger* if not mentioned otherwise. Numbers denote inter-river systems.

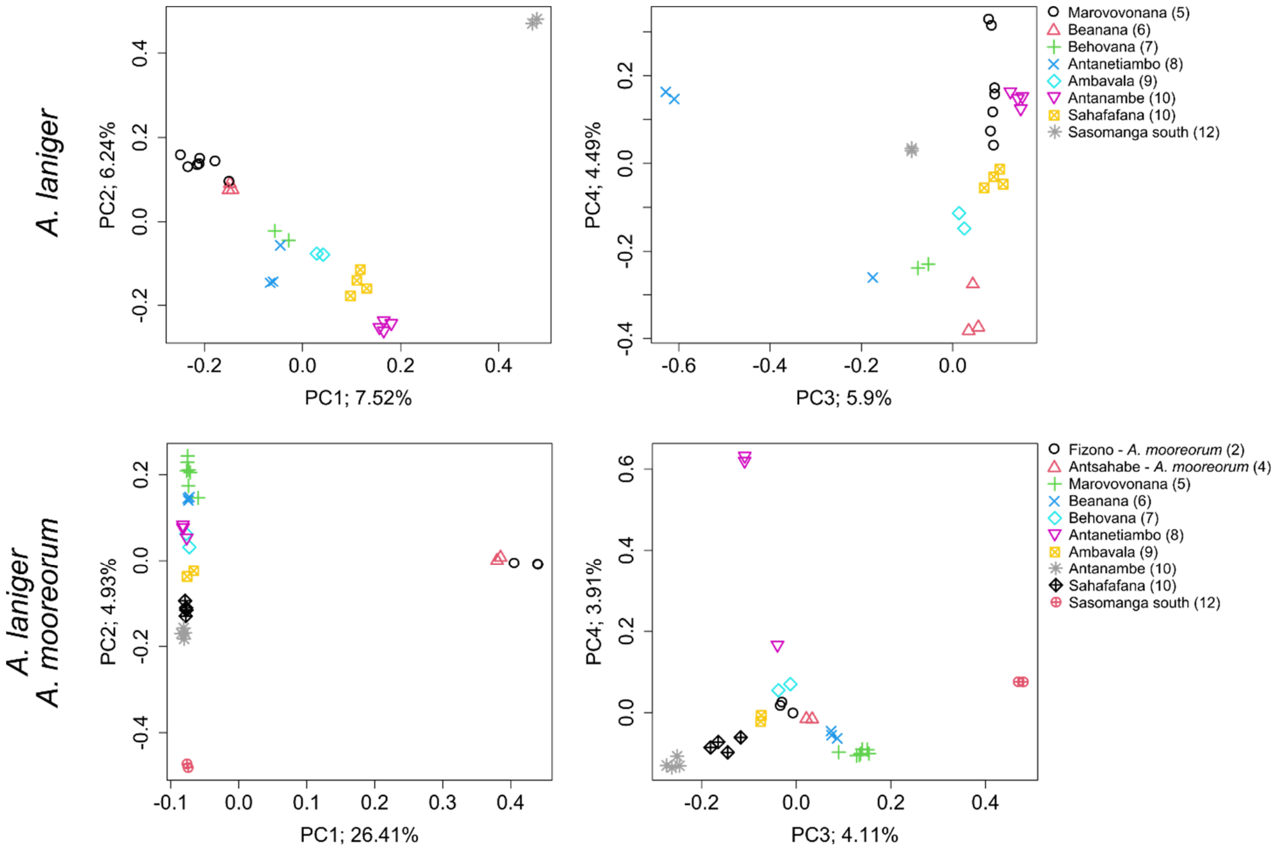


**Fig. S23:** Principal component analyses (PCA) of genetic data. Left column: PC1 plotted against PC2; Right column: PC3 plotted against PC4. Percentages indicate amount of variation explained by the PC. Population names refer to *A.laniger* if not mentioned otherwise. Numbers denote inter-river systems.

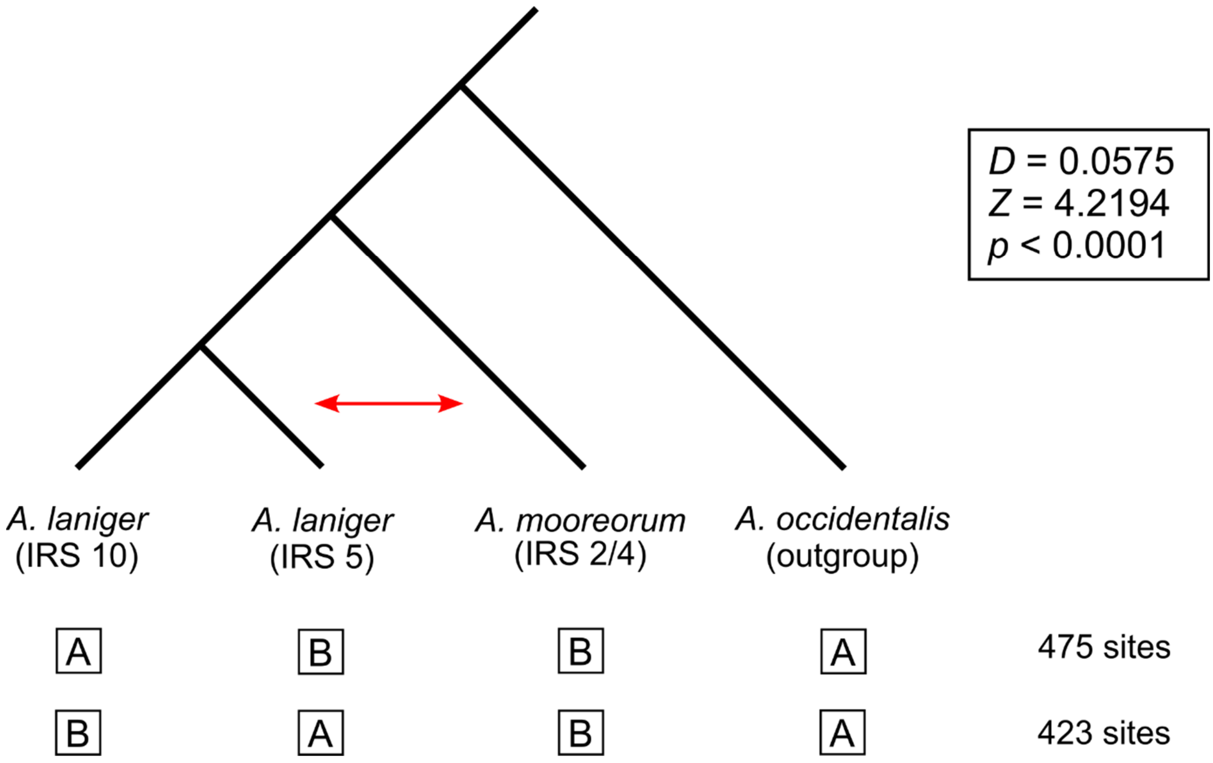


**Fig. S24:** Test for introgression between *A. laniger* (inter-river system/IRS 5) and the *A. mooreorum* populations from IRSs 2 and 4 (indicated by red arrow) via Patterson's *D* statistic. Significantly more shared sites were found between these clades (ABBA) than between *A. mooreorum* and the *A. laniger* populations from IRS 10 (BABA), providing evidence for introgression.

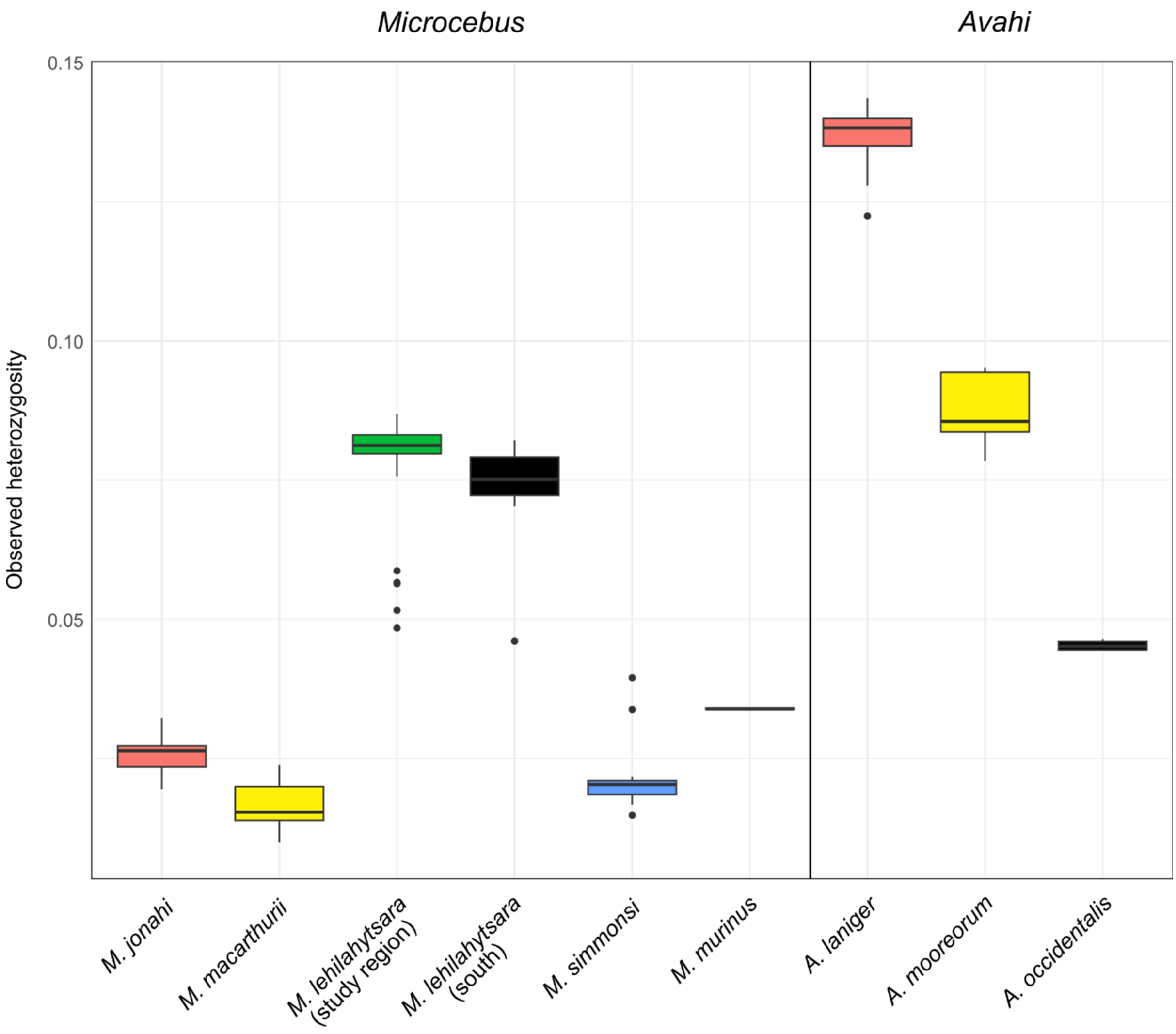


**Fig. S25:** Observed heterozygosity of *Microcebus* spp. and *Avahi* spp. individuals in the study region. Estimates for *M. lehilahytsara* populations south of the region as well as the *M. murinus* and *A. occidentalis* outgroups (black) were added for reference. Significance levels of pairwise comparisons estimated through Dunn's (*post hoc*) test after a Kruskal-Wallis test are given in Table S13. Sample sizes: $n_{M.\ jonahi} = 178$; $n_{M.\ macarthurii} = 15$; $n_{M.\ lehilahytsara\ (study\ region)} = 82$; $n_{M.\ lehilahytsara\ (south)} = 31$; $n_{M.\ simmonsi} = 36$; $n_{M.\ murinus} = 3$; $n_{A.\ laniger} = 27$; $n_{A.\ mooreroum} = 5$; $n_{A.\ occidentalis} = 4$.

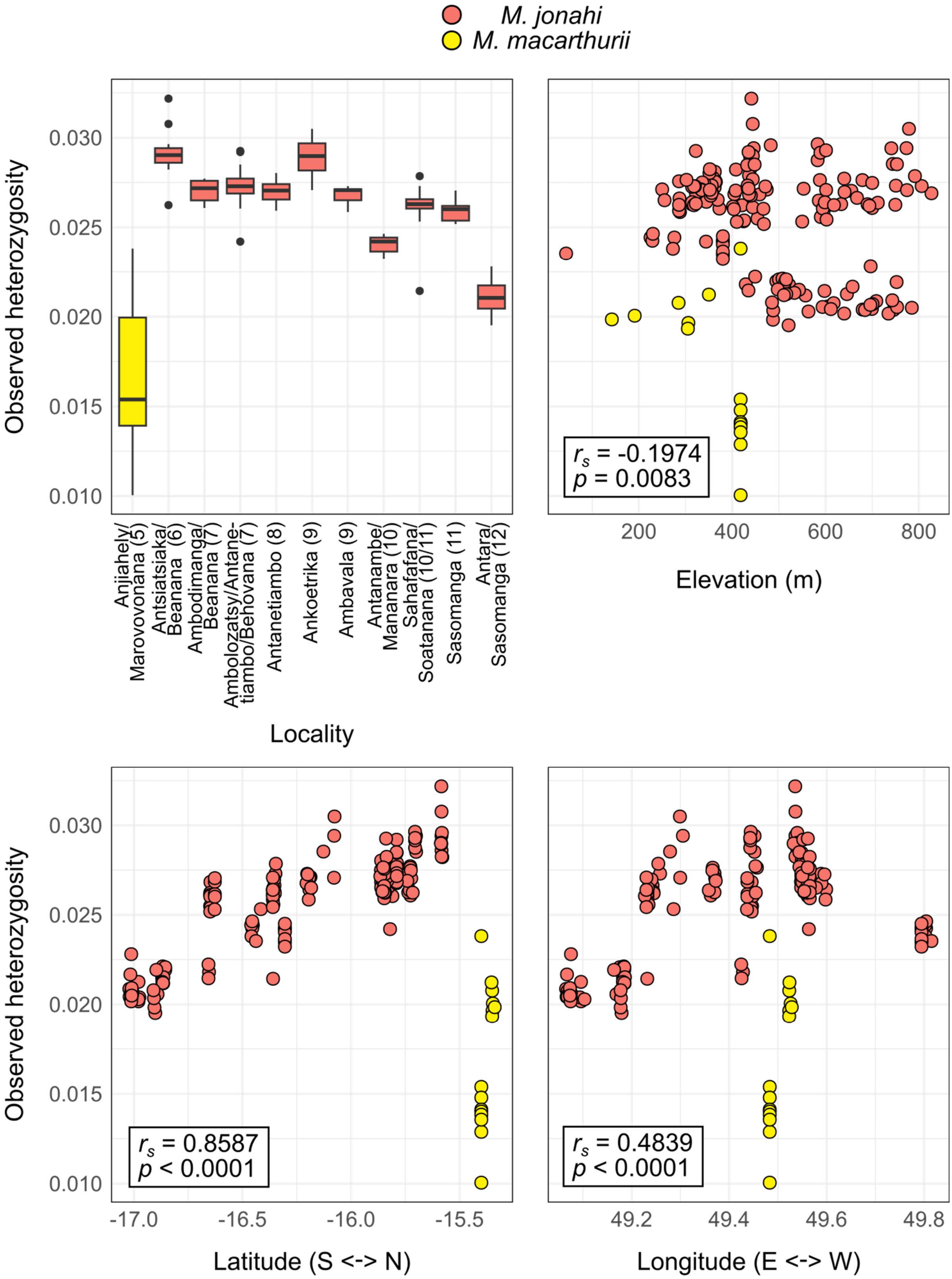


**Fig. S26:** Observed heterozygosity of *M. jonahi* and *M. macarthurii* individuals in the study region in northeastern Madagascar plotted per sampling locality (numbers denote inter-river system) and against elevation, latitude and longitude. Results of Spearman's rank correlation performed on *M. jonahi* data are given in the inlets. S: south; N: north; E: east; W: west. Sample sizes per population can be seen in Table S1.

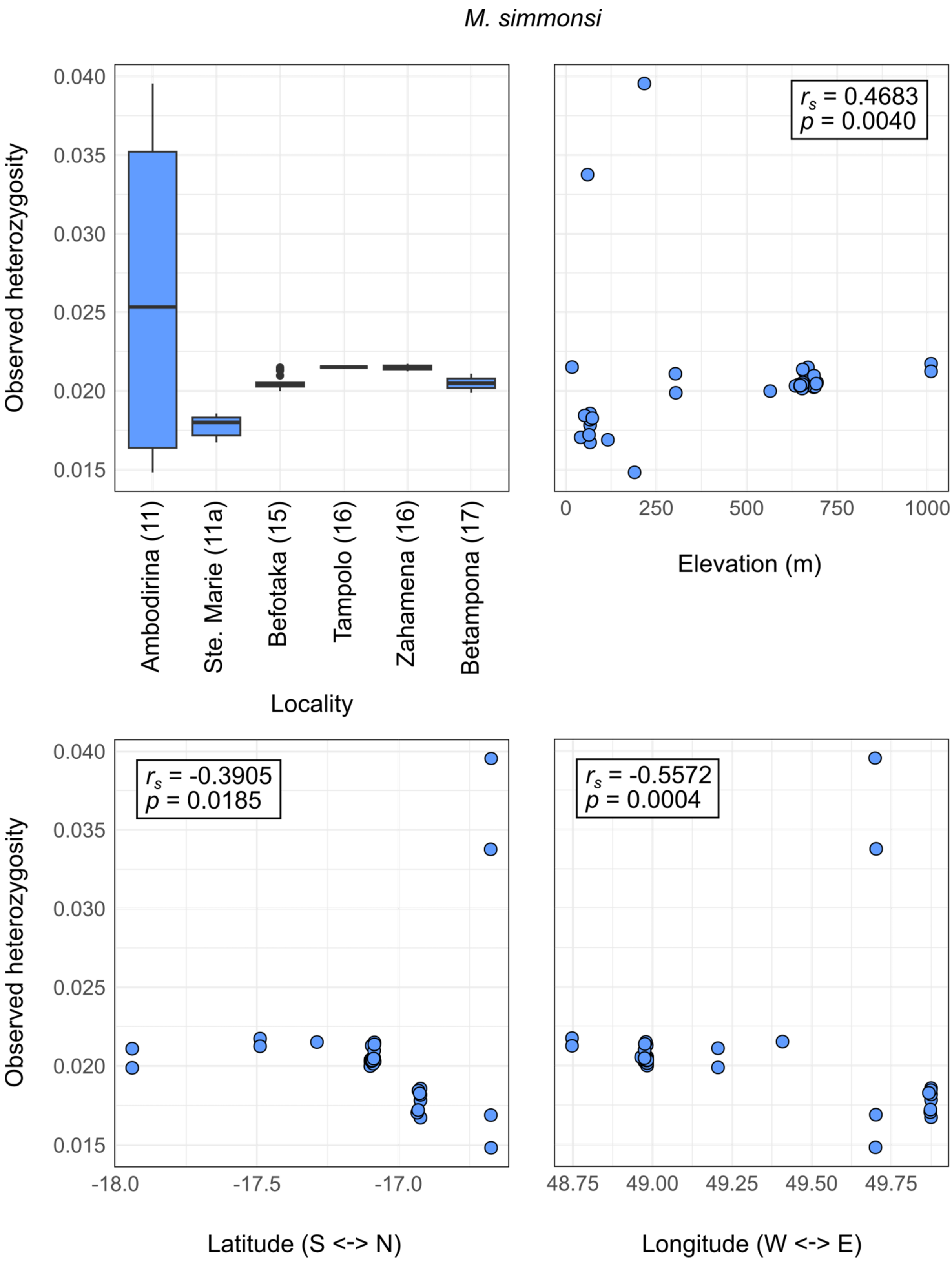


**Fig. S27:** Observed heterozygosity of *M. simmonsi* individuals in the study region in northeastern Madagascar plotted per sampling locality (numbers denote inter-river system) and against elevation, latitude and longitude. Results of Spearman's rank correlation are given in the inlets. S: south; N: north; E: east; W: west. Sample sizes per population can be seen in Table S1.

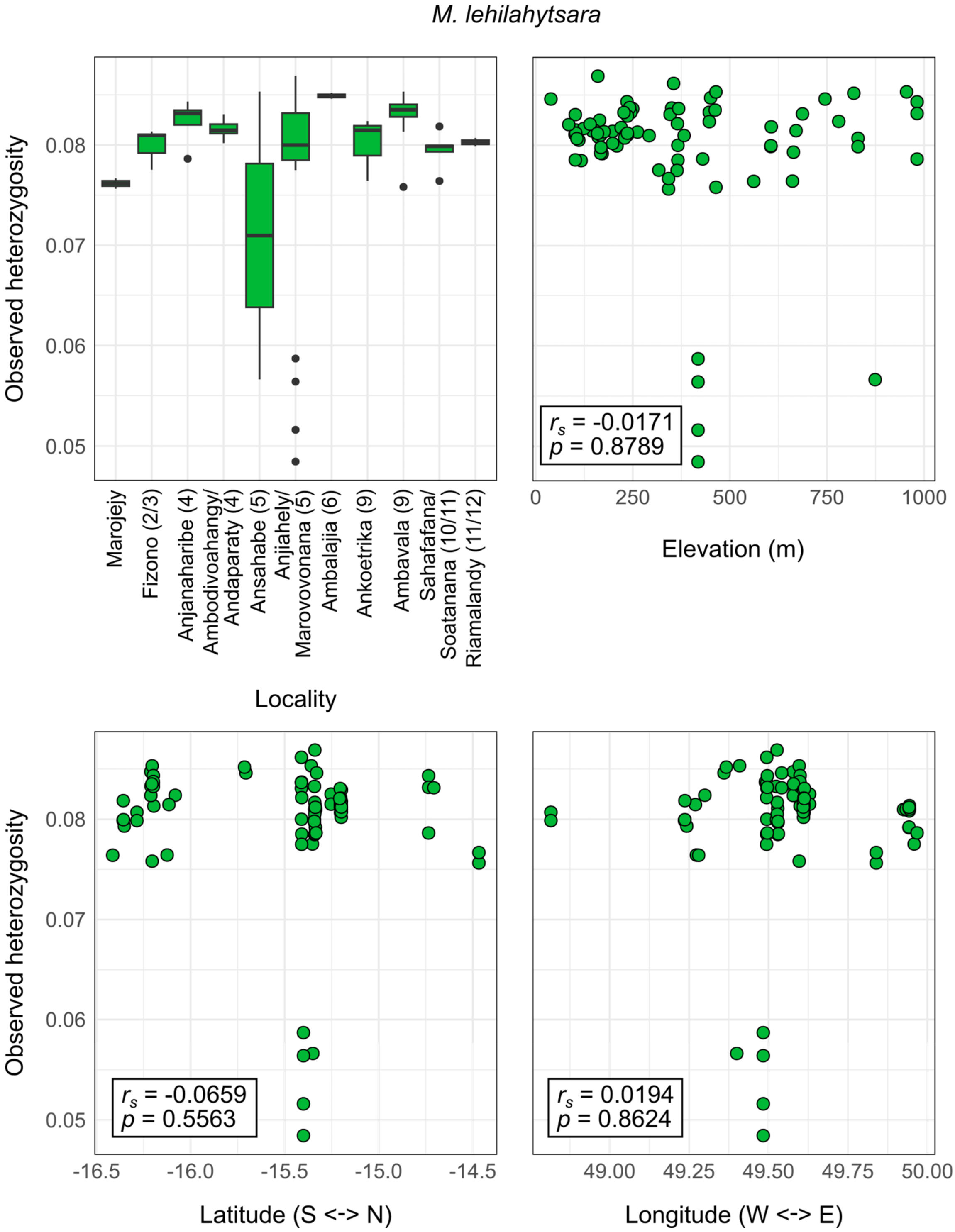


**Fig. S28:** Observed heterozygosity of *M. lehilahytsara* individuals in the study region in northeastern Madagascar plotted per sampling locality (numbers denote inter-river system) and against elevation, latitude and longitude. Results of Spearman's rank correlation are given in the inlets. The five outliers with observed heterozygosities below 0.06 are samples from Anjiahely (IRS5) with comparably high proportions of missing data (Tables S4.2 and S4.10). S: south; N: north; E: east; W: west. Sample sizes per population can be seen in Table S1.

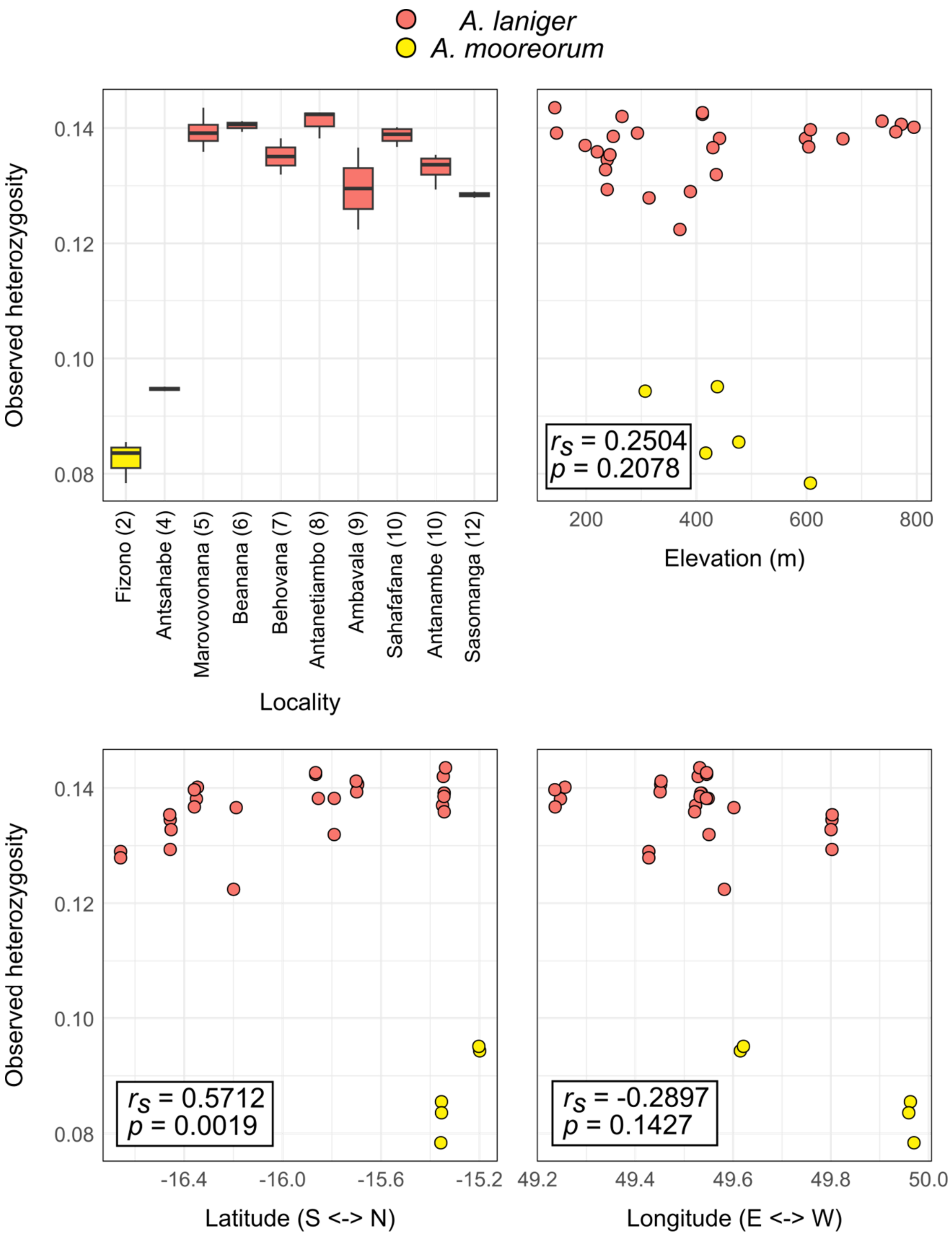


**Fig. S29:** Observed heterozygosity of *A. laniger* and *A. mooreorum* individuals in the study region in northeastern Madagascar plotted per sampling locality (numbers denote inter-river system) and against elevation, latitude and longitude. Results of Spearman's rank correlation performed on *A. laniger* data are given in the inlets. S: south; N: north; E: east; W: west. Sample sizes per population can be seen in Table S1.

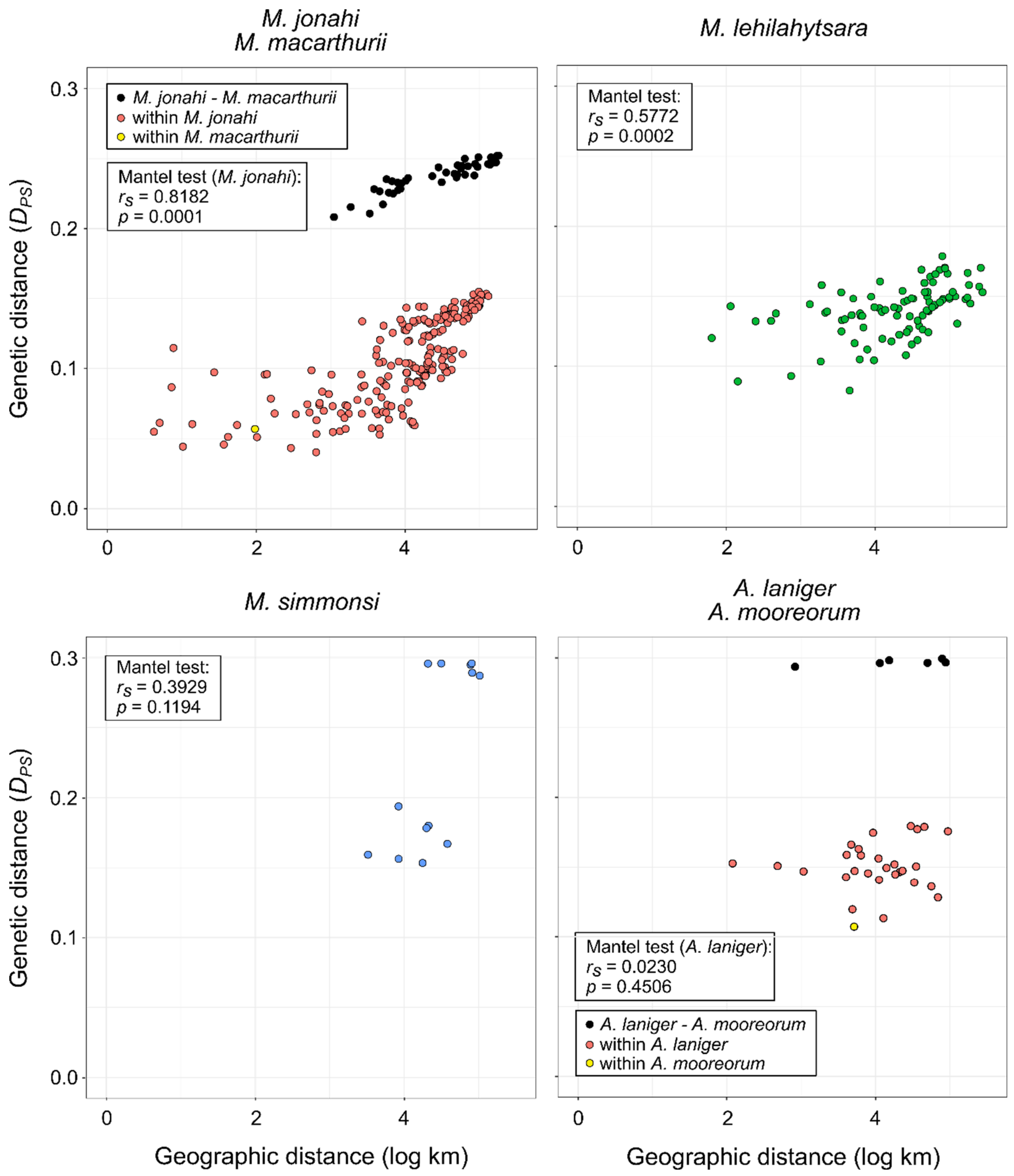


**Fig. S30:** Genetic distance $D_{PS}$ (1 – proportion of shared alleles) between populations of the study species plotted against geographic distance (on log scale). Geographic distance between populations was calculated as the mean of individual distances. Mantel tests based on Spearman's rank correlation $r_s$ were performed to test for isolation-by-distance. The tests for *M. jonahi* and *A. laniger* were conducted without *M. macarthurii* and *A. mooreorum* populations, respectively.

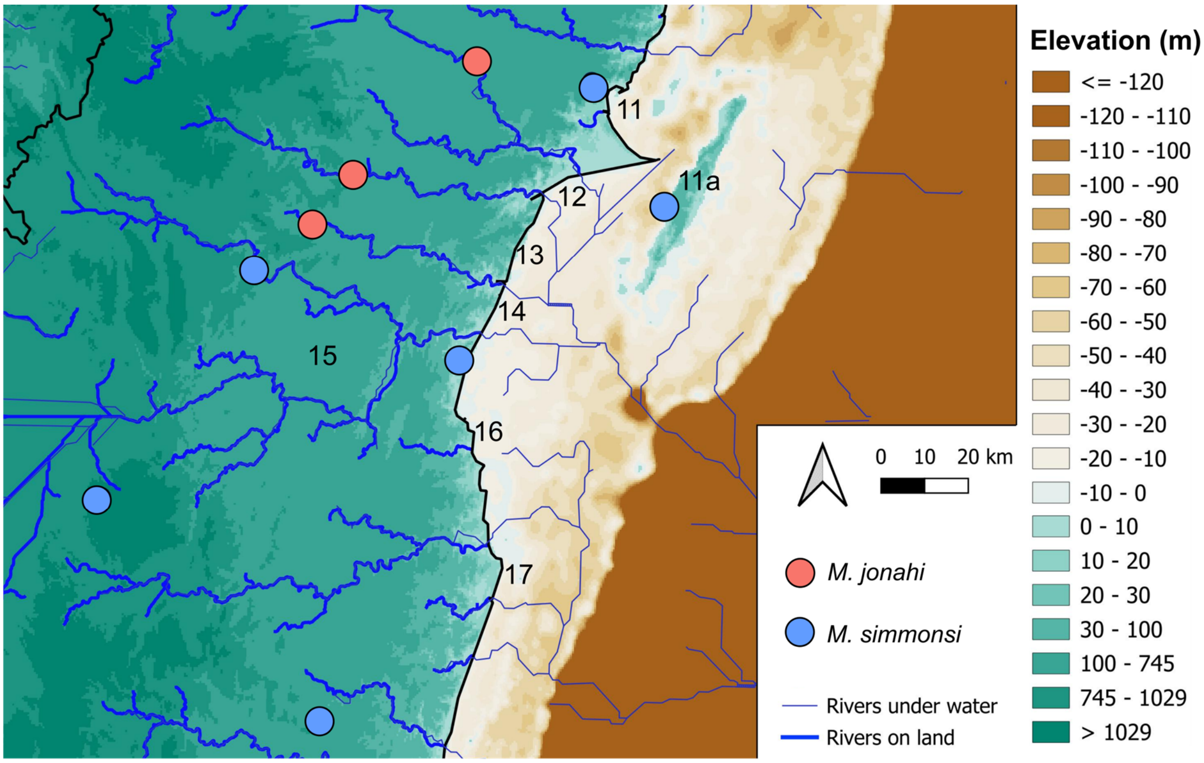


**Fig. S31:** River courses below sea level predicted via flow accumulation from a digital elevation model (Schüßler, 2025). According to the prediction, rivers separating inter-river systems (IRSs) 12 to 14 (indicated by numbers) converged into a single river when sea levels were lower than today (e.g., during the Last Glacial Maximum), potentially facilitating dispersal of *M. simmonsi* populations from the southern part of its distribution to Île Ste. Marie (IRS 11a). Colored dots indicate sites sampled in this study.
